# The Molecular Logic of Gtr1/2 and Pib2 Dependent TORC1 Regulation in Budding Yeast

**DOI:** 10.1101/2023.12.06.570342

**Authors:** Jacob H. Cecil, Cristina M. Padilla, Austin A. Lipinski, Paul R. Langlais, Xiangxia Luo, Andrew P. Capaldi

## Abstract

The Target of Rapamycin kinase Complex I (TORC1) regulates cell growth and metabolism in eukaryotes. Previous studies have shown that, in *Saccharomyces cerevisiae*, nitrogen and amino acid signals activate TORC1 via the highly conserved small GTPases, Gtr1/2, and the phosphatidylinositol 3-phosphate binding protein, Pib2. However, it was unclear if/how Gtr1/2 and Pib2 cooperate to control TORC1. Here we report that this dual regulator system pushes TORC1 into at least three distinct signaling states: (i) a Gtr1/2 on, Pib2 on, rapid growth state in nutrient replete conditions; (ii) a Gtr1/2 inhibited, Pib2 on, adaptive/slow growth state in poor-quality growth medium; and (iii) a Gtr1/2 off, Pib2 off, quiescent state in starvation conditions. We suggest that other signaling pathways work in a similar way, to drive a multi-level response via a single kinase, but the behavior has been overlooked since most studies follow signaling to a single reporter protein.

## INTRODUCTION

The Target of Rapamycin kinase Complex 1 (TORC1) is the master regulator of cell growth and metabolism in eukaryotes^1–3^. In the presence of pro-growth hormones and abundant nutrients, TORC1 is active and drives growth by stimulating protein, ribosome, lipid, and nucleotide synthesis^1–9^. In contrast, when hormone or nutrient levels drop, TORC1 is inhibited, causing the cell to switch from anabolic to catabolic metabolism and eventually enter a quiescent state^10–12^.

Numerous proteins and pathways have been shown to regulate TORC1, including: (i) the small GTPases Rag A/B and C/D^13–19^ and the associated GTPase Activation Protein (GAP), GATOR1/2^20^; (ii) the small GTPase Rheb and its associated GAP, the Tuberous Sclerosis Complex^21–23^; (iii) the AMP activated protein kinase^24–26^; (iv) the Nemo-like kinase^27^; (v) the cAMP-dependent protein kinase^28^; and (vi) the small GTPase ADP-ribosylation factor 1^29,30^. However, it remains unclear how the proteins/pathways listed above (and others) cooperate to control TORC1.

Here, to address this question, we examine signal integration at TORC1 in the simple model organism, *Saccharomyces cerevisiae*.

Previous work in *S. cerevisiae* has shown that:

(1) Amino acid and nitrogen signals are transmitted to TORC1 via a heterodimeric pair of small GTPases that are homologous to RagA/B and RagC/D, called Gtr1 and Gtr2^18,19^. Specifically, when cells are grown in medium containing a high concentration of amino acids (or a high-quality nitrogen source), Gtr1/2 are in their GTP and GDP bound forms, respectively, and bind to/activate TORC1^18,19^. However, once amino acid/nitrogen levels fall, SEAC (a homolog of GATOR1/2) triggers GTP hydrolysis at Gtr1 to drive the complex into the Gtr1-GDP, Gtr2-GTP bound state, and inhibit TORC1^31–35^.
(2) TORC1 is also regulated by the phosphatidylinositol 3-phosphate binding protein, Pib2^36,37^. Much less is known about Pib2 than Gtr1/2, but several facts are clear: First, Pib2 binds directly to TORC1 and activates the complex via its highly conserved C-terminal domain (CAD) domain^37–41^. Second, Pib2 can repress TORC1 via its N-terminal inhibitory domain (NID)^41^. Third, Pib2 activates TORC1 in response to glutamine^40,42,43^.

But how and why do Gtr1/2 and Pib2 work together to regulate TORC1?

Most data suggest that Gtr1/2 and Pib2 act in parallel (redundantly) to activate TORC1^36^. For example, yeast missing either Gtr1/2 or Pib2 grow well in nutrient rich media, while *gtr1*/*2*τι*pib2*τι cells are sick/dead (a phenotype that can be rescued by a hyperactive TOR allele)^37^. Furthermore, *gtr1/2*τι and *pib2*τι cells have strong TORC1 activity as measured by the downstream reporters phospho-Rps6 and Sch9, while transient repression of Pib2 in a *gtr1/2*τι cell line blocks TORC1 signaling to these same proteins^37,42^.

However, other data suggest that the impact that Gtr1/2 and Pib2 have on TORC1 signaling is more complex: First, *gtr1*τι, *gtr2*τι, *gtr1/2*τι and *pib2*τι cells are all hypersensitive to the TORC1 inhibitor rapamycin^41^. Second, both *gtr1/2*τι and *pib2*τι cells fail to activate TORC1 when leucine or glutamine are added back to cells treated with rapamycin^44^. Third, phosphorylation of the TORC1 target Npr1, and its substrate Par32, are sensitive to deletion of either Pib2 or Gtr1/2^37,45^.

In this report, we build on the previous studies by examining the impact that Gtr1/2 and Pib2 have on TORC1 signaling across the proteome, using a combination of phosphoproteomics and standard reporter assays. The resulting data show that Gtr1/2 and Pib2 are both required for full TORC1 activation. Importantly, however, deletion of Gtr1/2 or Pib2 only blocks signaling to a subset of the TORC1 substrates—primarily those involved in amino acid metabolism and nutrient transport. These observations lead us to propose a new model where partial starvation triggers metabolic reprograming via TORC1 (by inactivating Gtr1/2 or Pib2) but does not block cell growth. And in follow up experiments, we confirm that this is, indeed, the case. Specifically, we show that when yeast are first transferred from medium containing a high-quality nitrogen source, to medium containing a low-quality nitrogen source, TORC1 is completely inhibited to block growth and activate metabolic reprogramming. Then, as the cells adapt to the low-quality nitrogen source, Pib2 is turned on again to re-initiate growth, while Gtr1/2 remains inhibited, or partially inhibited, to ensure that the cells continue to activate the metabolic pathways and transporters necessary for adaptation/survival.

Thus, the TORC1 circuit in yeast uses two different amino acid/nitrogen signaling proteins to drive the cell into a rapid growth state, an adaptive growth state, or a quiescent state, depending on the environmental conditions. We argue that other signaling pathways probably work in a similar way, but multi-level responses are overlooked since most studies follow signaling to a single reporter.

## RESULTS

### Reporter (pRps6) based analysis of Gtr1/2 and Pib2 dependent signaling

Gtr1/2 and Pib2 are thought to transmit leucine, glutamine, and other amino acid signals to TORC1^36^. To test this model and learn more about the cooperation between Gtr1/2 and Pib2, we followed the phosphorylation of a downstream reporter of TORC1 activity (Rps6) in wild-type, *gtr1*τι, and *pib2*τι cells during amino acid starvation^46,47^. These experiments showed that all three strains have the same TORC1 activity in nutrient replete medium (time 0, Fig. 1), consistent with the idea that Gtr1/2 and Pib2 work in parallel (redundantly) to activate TORC1. These experiments also showed that TORC1 turns off efficiently in the absence of Gtr1 or Pib2 during leucine and complete amino acid starvation (Fig. 1). However, the results in glutamine starvation stood out; TORC1 remained active in the wild-type strain, and partially active in the *pib2*τι strain, but was rapidly inhibited in the *gtr1*τι strain (Fig. 1).

**Figure 1.**
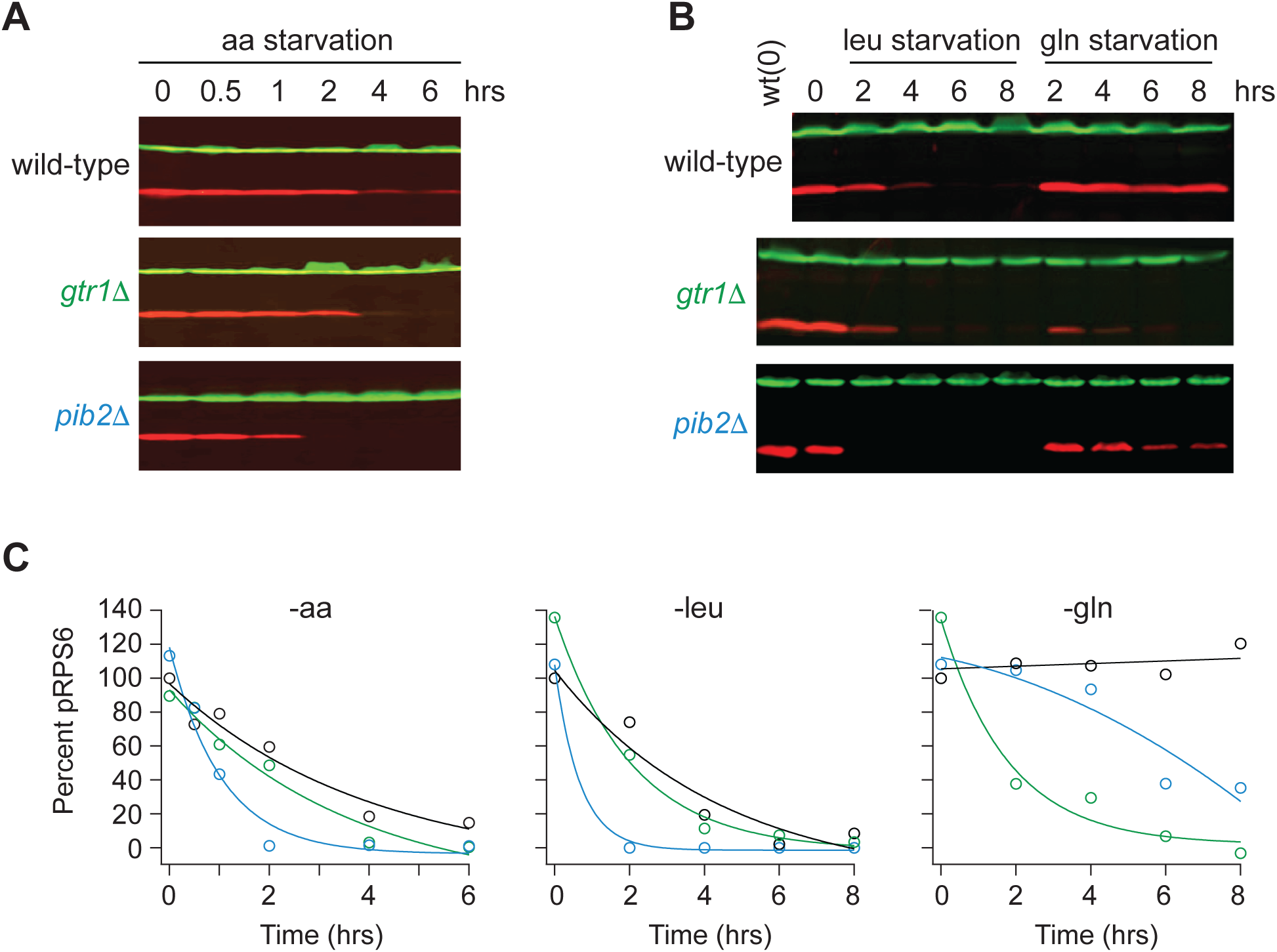
Impact of Gtr1/2 and Pib2 on TORC1 signaling during amino acid starvation. **(A)** TORC1 activity measured before and after complete amino acid starvation in wild-type, *gtr1Δ*, and *pib2Δ* strains using a Western blot with anti phospho-Rps6 (red) and anti PGK (green) antibodies. **(B)** TORC1 activity measured as in (A) but during leucine and glutamine starvation. Glutamine starvation was triggered by transferring the cells from synthetic complete medium (SC), to SC medium missing glutamine and containing 2 mM methionine sulfoximine (MSX; a glutamine synthetase inhibitor). Leucine starvation was triggered by transferring the three leu^-^ strains from SC medium to SC medium missing leucine. **(C)** Values showing the ratio of the p-Rps6 signal divided by the PGK (loading control) signal in each lane from (A) and (B), relative to the value for the wild-type strain at time = 0.

To learn more about the interaction between Gtr1/2 and Pib2 during glutamine starvation, we followed Rps6 phosphorylation in cells missing the N-terminal inhibitory domain of Pib2 (*pib2NID*τι). The *pib2NID*τι mutation completely blocked TORC1 inhibition in the *gtr1*τι background but had no impact on TORC1 signaling in a wild-type background (Fig. 2a, b), demonstrating that Pib2 helps drive TORC1 inactivation during glutamine starvation (at least in the absence of Gtr1).

**Figure 2.**
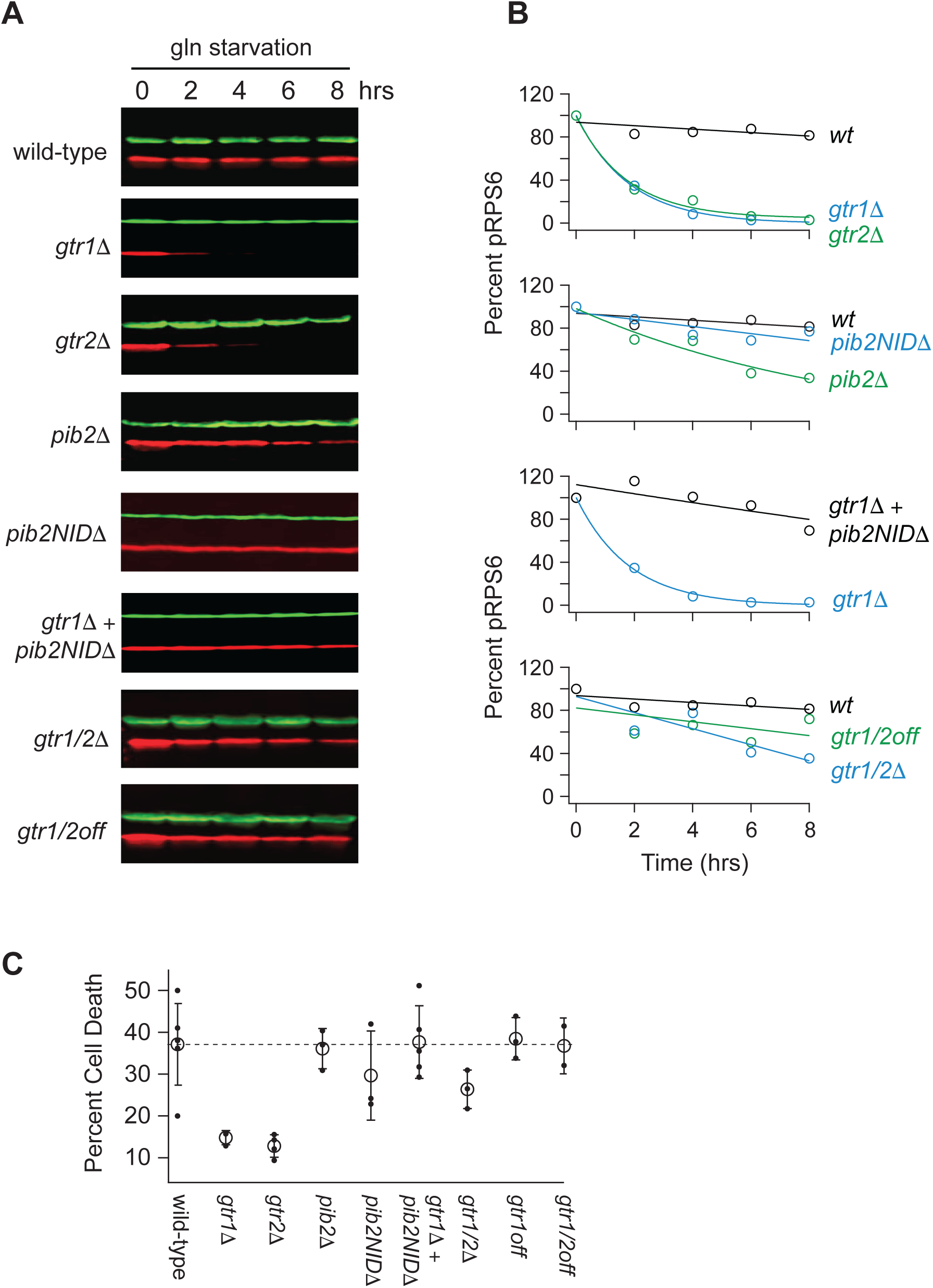
Impact of Gtr1/2 and Pib2 on TORC1 signaling during glutamine starvation. **(A)** TORC1 activity measured during glutamine starvation in *gtr1Δ*, *gtr2Δ*, *pib2Δ, pib2NIDΔ, gtr1Δpib2NIDΔ, gtr1Δgtr2Δ,* and *gtr1/2^off^*strains using a Western blot, as described in Figure 1. **(B)** Values showing the ratio of the p-Rps6 signal divided by the PGK (loading control) signal in each lane from (A) relative to the value for the wild-type strain at time = 0. **(C)** Fraction of cells that are dead after six hours of glutamine starvation for each strain listed in (B) as measured using SYTOX green labelling and a fluorescence microscope. The open circles and error bars show the average and standard deviation from four replicates (filled circles), with >200 cells analyzed per replicate, per strain.

We then measured Rps6 phosphorylation in a strain missing both Gtr1 and Gtr2 (*gtr1*τι*gtr2*τι), and a strain carrying mutations that lock Gtr1 and 2 into their inactive Gtr1-GDP and Gtr2-GTP bound forms (Gtr1^S20L^ and Gtr2^Q66L^; Gtr1/2^off^ for short^18^). To our surprise, TORC1 remained active in both strains (Fig. 2a, b), demonstrating that Pib2 can only inhibit TORC1 during glutamine starvation in strains where the Gtr1/2 complex is partially disrupted (*gtr1*τι and *gtr2*τι strains) but not completely absent or inactive (Fig. 2a, b).

We also saw the same pattern when we examined cell survival in glutamine starvation. Wild-type, *pib2*τι, *pib2NID*τι, *pib2NID*τι*gtr1*τι, Gtr1/2^off^, and the other cell lines that failed to inactivate TORC1, all die at a higher rate during glutamine starvation than the *gtr1*τι and *gtr2*τι cells--presumably because they do not arrest cell growth in a timely manner (Fig. 2c).

In sum, most of our measurements of Rps6 signaling and cell death fit with the prevailing model of TORC1 regulation, where Gtr1/2 and Pib2 (i) act in parallel (redundantly) to activate TORC1 in nutrient replete conditions, and (ii) are switched off during amino acid starvation to inhibit TORC1. However, we were left with a puzzle. Our data also revealed that Pib2 can transmit glutamine starvation signals to TORC1, but those signals have little to no impact on TORC1 activity in the presence of an intact Gtr1/2 complex. Why would this be? We hypothesized that signals transmitted through Pib2 alone, or Gtr1/2 alone, do impact TORC1 signaling in a wild-type background, but the response is just not apparent at the level of Rps6 phosphorylation.

### Proteome wide analysis of Gtr1/2 and Pib2 dependent signaling

To learn more about Gtr1/2 and Pib2 signaling, we used mass spectrometry to quantify the global protein phosphorylation levels in wild-type, *gtr1*τι*gtr2*τι, and *pib2*τι cells grown in nutrient replete medium (SC), and in wild-type cells treated with the TORC1 inhibitor rapamycin (all in quadruplicate). In the end, we were able to quantify the level of 7807 phosphopeptides (covering 5325 phosphosites on 1686 proteins) across the 16 samples (Table S1). 445 of these phosphopeptides (covering 362 phosphosites on 301 proteins) were up- or down-regulated in response to rapamycin treatment, deletion of Gtr1/2, and/or the deletion of Pib2 (>2-fold change and p<0.01 in one or more strain/condition). More specifically, 175 phosphopeptides were down-regulated in rapamycin (Figs. 3 and 4), 187 phosphopeptides were up-regulated in rapamycin (Fig. 4S1), and 83 phosphopeptides were up or down regulated in the *gtr1*τι*gtr2*τι or *pib2*τι cells, but not in the rapamycin treated cells (Fig. 4S2).

**Figure 3.**
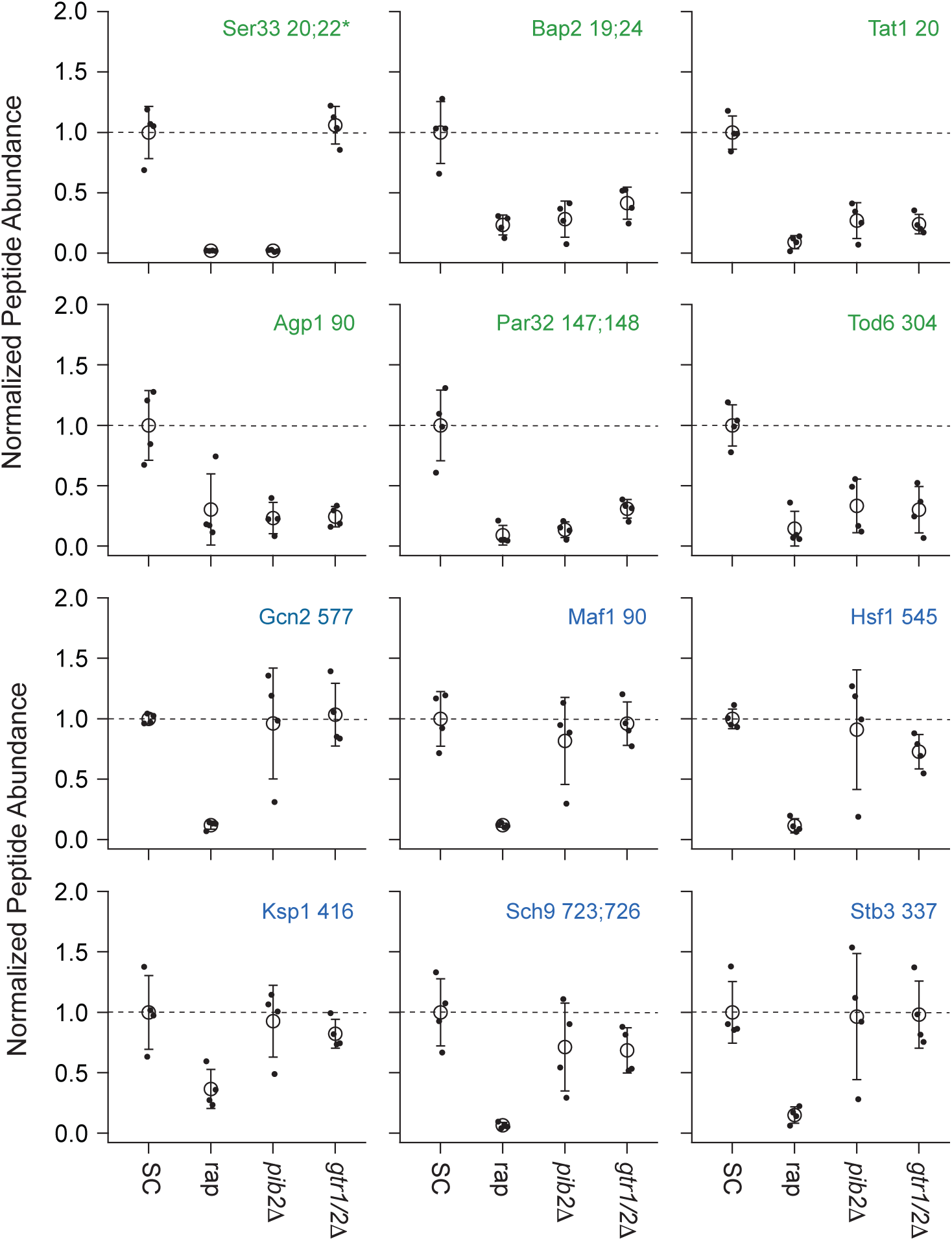
Impact of rapamycin, Gtr1/2 deletion, and Pib2 deletion on select TORC1-dependent phosphopeptides. Normalized peptide abundance for select phosphopeptides extracted from wild type cells growing in SC medium, wild-type cells growing in SC medium and treated with 200 nM rapamycin for 30 min, or *pib2Δ* or *gtr1Δgtr2Δ* cells growing in SC medium (as labeled). The open circles and error bars show the average and standard deviation from four replicates (filled circles) and all data is divided by the average signal (for the relevant peptide) in the wild-type strain growing in SC medium. The phosphopeptides are named based on the protein they are from, followed by the number of the phosphorylated residue(s). Entries with an asterisk (*) indicate that there are other possible phosphorylation site assignments (see Table S1). Phosphopeptides labeled in green (top panels) change significantly in the *pib2Δ* and/or *gtr1Δgtr2Δ* backgrounds.

**Figure 4.**
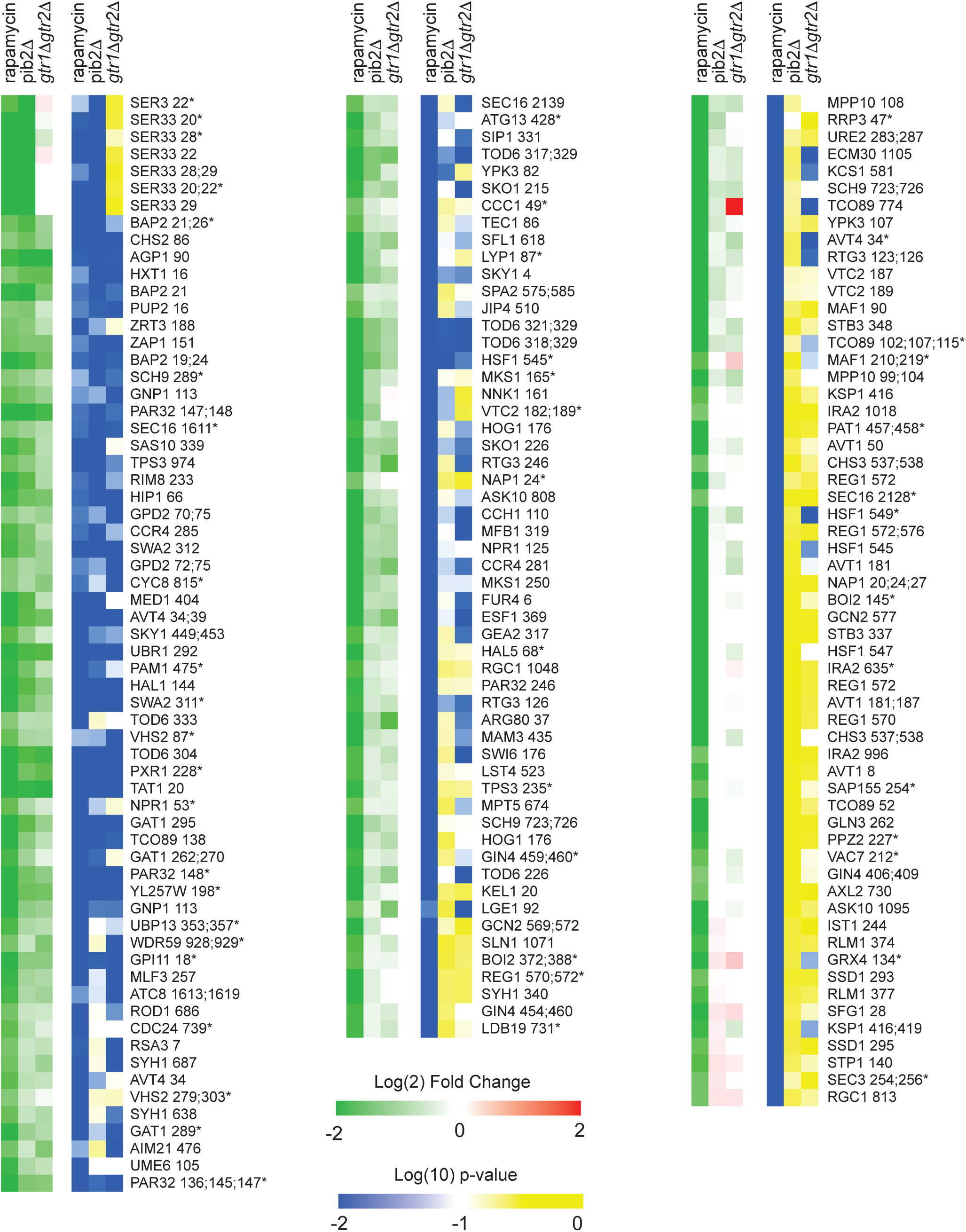
Phosphopeptides significantly downregulated in rapamycin. Heatmap showing the abundance of all the phosphopeptides that are downregulated in rapamycin (>2-fold with p-value <0.01). Columns 1-3 (green-red) show the peptide levels after 30 min rapamycin treatment, in the *pib2Δ* strain growing in SC medium, and in the *gtr1Δgtr2Δ* strain growing in SC medium (as labelled), all compared to those in the wild-type strain growing in SC medium. The values are the average from four replicate experiments. Columns 4-6 (blue-yellow) show the statistical significance of any change in columns 1-3 based on a t-test. The expression data is ordered based on the fraction of the rapamycin response found in the *pib2*11 strain (top left, to bottom right). The phosphopeptides are named based on the protein they are from, followed by the number of the phosphorylated residue(s). Entries with an asterisk (*) indicate that there are other possible phosphorylation site assignments (see Table S1).

**Figure 4 S1.**
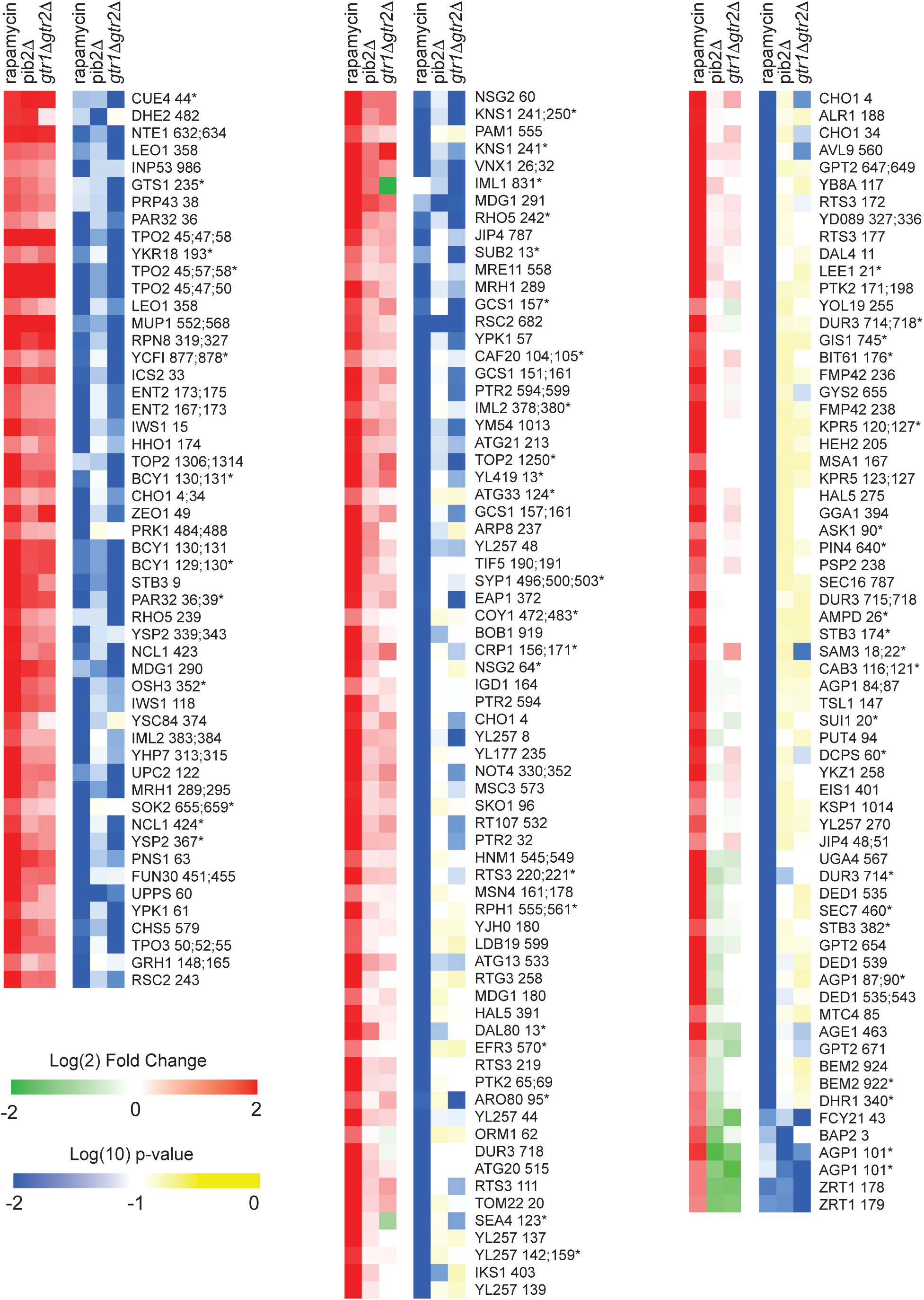
Phosphopeptides significantly upregulated in rapamycin. Heatmap showing the abundance of all the phosphopeptides that are upregulated in rapamycin (>2-fold with p-value <0.01). The figure is laid out as described for Fig. 4.

**Figure 4 S2:**
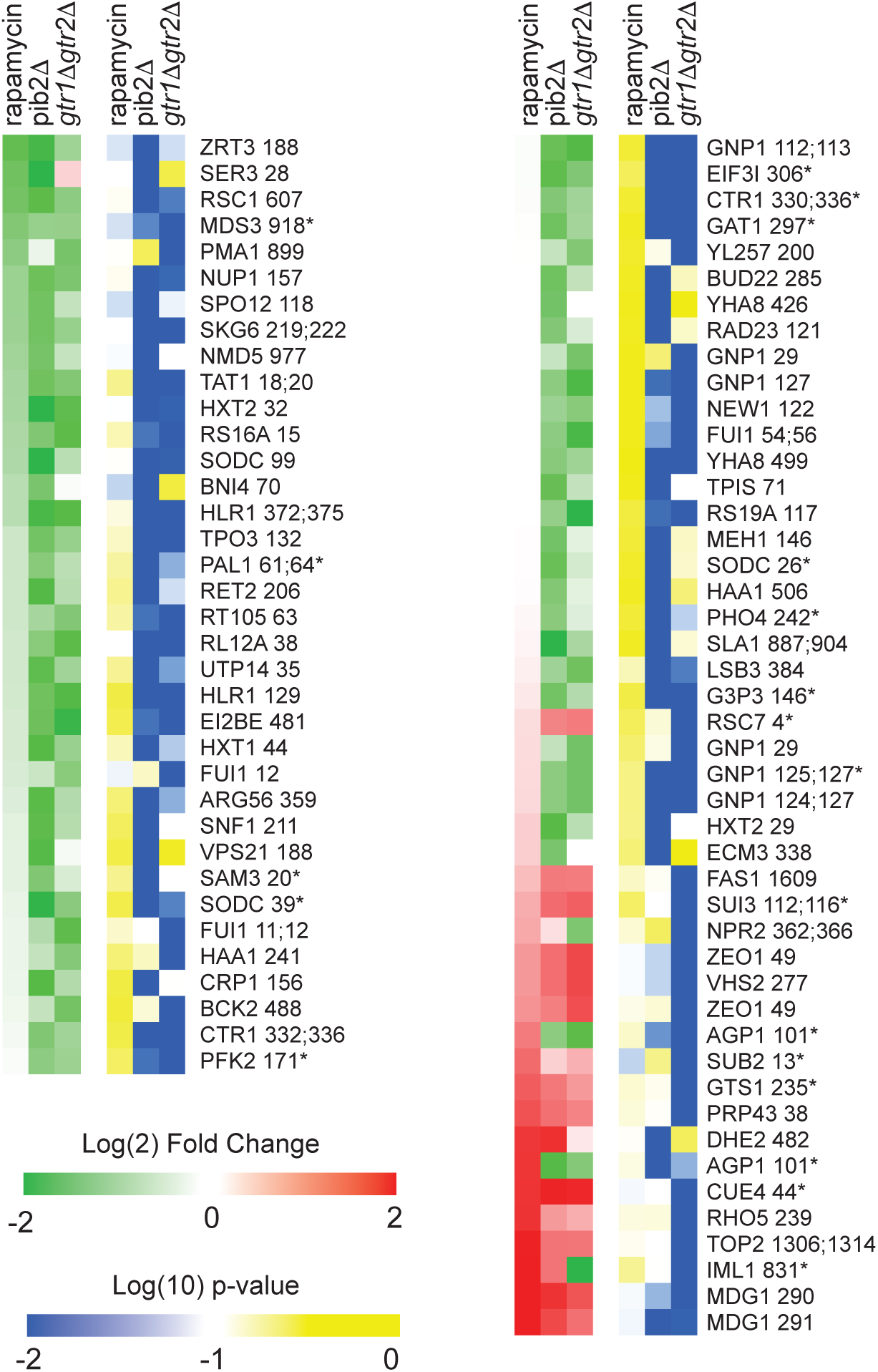
Phosphopeptides significantly up/downregulated by deletion Gtr1/2 or Pib2 but not by rapamycin. Heatmap showing the abundance of the phosphopeptides that are up or down regulated in the Gtr1/2 and/or Pib2 delete strains (>2-fold with p-value <0.01) but with no statistically significant change in rapamycin treatment (i.e., not assigned to Fig. 4, or 4S1). The figure is laid out as described for Fig. 4. Note that a significant number of the peptides in this heatmap are sensitive to rapamycin but failed to make the (2-fold and p<0.01) cutoff threshold due to noise (see top left column, and bottom right column). Some of the other data in this heatmap may also represent noise—particularly for the peptides where there is a significant change in the *pib2Δ* or *gtr1Δgtr2Δ* strains, but not both. Where there is a *bona fide* change in the *pib2Δ* and *gtr1Δgtr2Δ* strains but no change in rapamycin, for example at the multiple target sites in the glutamine transporter Gnp1 (top and middle, right column), we suspect that there are competing TORC1 dependent regulatory events. The first is a Gtr1/2 and Pib2 sensitive phosphorylation event. The second is a TORC1 repressed (Gtr1/2 and Pib2 insensitive) phosphorylation event. This dual control mechanism could be used to activate/repress proteins during intermediate, but not complete, starvation.

We focused our analysis on the 175 phosphopeptides that are downregulated in rapamycin since they cover most of the well-known targets of TORC1 and its downstream effectors, including Sch9, Tod6, Maf1, Stb3, Ypk3, Atg13, Mks1, Nnk1, Npr1, Par32, Avt1, Avt4, Sky1, Gat1, Gln3, Ume6, Rtg3, Lst4, Gcn2, Tco89, Ssd1, and Stp1 (Figs. 3 and 4)^3,48–55^. This dataset revealed two novel aspects of Gtr1/2 and Pib2 signaling. First, TORC1 drives the phosphorylation of several residues near the N-termini of Ser3 and Ser33 (Figs. 3, 4), homologous 3-phosphoglycerate dehydrogenases that catalyze the first step in serine and glycine synthesis^56^. Remarkably, these phosphorylation reactions are completely dependent on Pib2, but are not altered in the *gtr1*τι*gtr2*τι strain (Figs. 3, 4). Second, many TORC1 dependent phosphorylation events (outside of Ser3/33) depend heavily on *both* Pib2 and Gtr1/2 (green labels, Fig. 3, and left columns, Figs. 4 and 4S1). However, other phosphorylation events are unperturbed by the deletion of Gtr1/2 or Pib2 (blue labels, Fig. 3 and right columns, Fig. 4 and 4S1). In fact, there is a strong correlation between the impact that Pib2 and Gtr1/2 have on the TORC1 dependent phosphorylation (R=0.70 excluding Ser3/33; Fig. 4), but that impact runs the gamut from matching the impact of rapamycin, to zero (top left, to bottom right, continuum; Fig. 4 and 4S1).

Examining the list of proteins that are heavily dependent on Pib2 and Gtr1/2, we noticed an obvious trend; 8/11 of the proteins with the strongest reliance on Pib2 and Gtr1/2 are involved in nutrient transport and utilization (top left column, Fig. 4). These proteins include the amino acid transporters Bap2, Agp1, Gnp1 and Hip1^57–60^; the zinc transporter Zrt3 and the Zinc-regulated transcription factor, Zap1^61,62^; the hexose transporter Hxt1^63^; and the TORC1 dependent regulator of amino acid transporter activity, Par32^64^. This trend was also clear when we carried out GO analysis on the full list of Gtr1/2 and Pib2 dependent proteins (left column, Fig. 4), which includes the key TORC1 dependent regulator of amino acid metabolism, Npr1^64–66^, and the amino acid transporter Tat1^67^ (amino acid and aromatic amino acid transporter activity were the top hits; p=1e-4).

In contrast, the proteins that are dependent on TORC1, but not Gtr1/2 or Pib2 (right column, Figure 4), tend to be involved in cell signaling (p=1e-4) and the regulation of cell growth (biosynthetic processes, p=9e-3). This group of proteins includes many of the key regulators of ribosome and protein synthesis, including Sch9, Maf1, Stb3, Gcn2, and Kcs1^6,51,52,55,68^, as well as the stress response factor Hsf1^69^ and the autophagy regulator Ksp1^70^ (Figs. 3 and right column, Fig. 4).

Thus, the view of Gtr1/2 and Pib2 signaling that we and others built up by following Rps6 phosphorylation is misleading (Figs. 1 and 2). Gtr1/2 and Pib2 do not act redundantly during steady state growth, but instead, are both required for full TORC1 activation. It is just that some TORC1 pathway targets (like Rps6) are efficiently phosphorylated even when TORC1 is partially active.

### TORC1 and Pib2 dependent regulation of Ser33

To gain insight into the function of, and mechanism underlying, TORC1 and Pib2 dependent signaling to Ser33 (the dominant enzyme in the Ser3/33 pair^71^) we first set out map the TORC1 dependent phosphorylation sites on Ser33. Our global phosphoproteomics experiments had already identified serine 20, 22, 28, and 29 as TORC1 and Pib2 dependent sites (Figs. 3 and 4). However, to see if we missed any sites due to under-sampling, we also immunopurified Ser33-FLAG from cells grown in SD medium +/- rapamycin and mapped the phosphorylation sites using mass spectrometry. These experiments (and previously published phosphoproteomics data^51^) indicated that serine 27, serine 33, and threonine 31 are also TORC1 dependent sites (Fig. 5a; Table S2).

**Figure 5.**
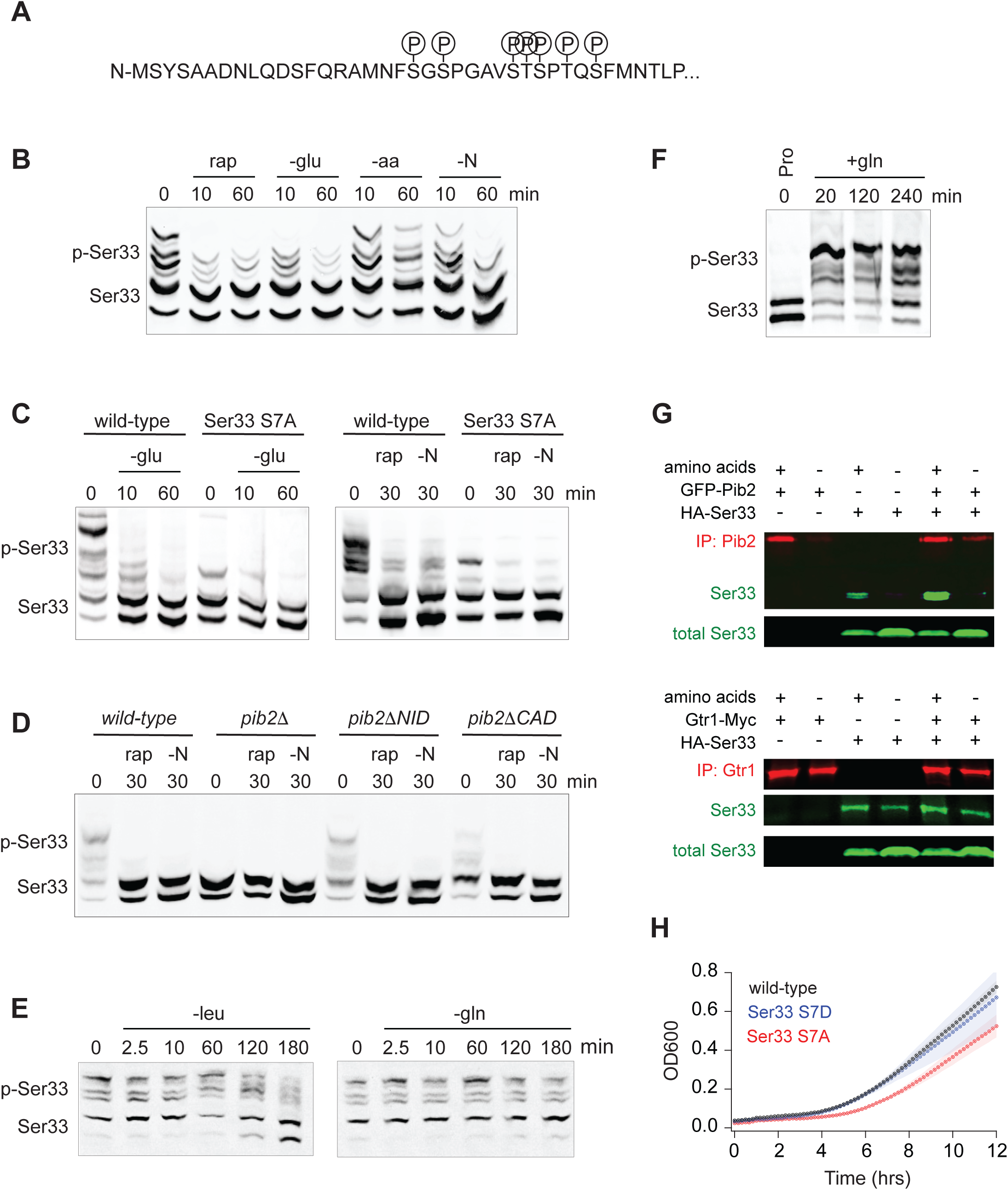
Function of, and mechanism underlying, the TORC1 and Pib2 dependent phosphorylation of the phosphoglycerate dehydrogenase Ser33. **(A)** TORC1-dependent phosphorylation sites compiled from global phosphoproteomics data, and mass-spectrometry based analysis of Ser33 immunopurified from cells in mid-log phase before and after treatment with 200 nM rapamycin. **(B)** Phos-tag gel measuring Ser33 phosphorylation before and after treatment with 200 nM rapamycin (rap) or starvation for glucose (-glu), amino acids (-aa), or all nitrogen (-N). **(C)** Phos-tag gel comparing Ser33 phosphorylation in wildtype and Ser33^S7A^ cells following glucose starvation, rapamycin treatment, or nitrogen starvation. (**D**) Phos-tag gel examining Ser33 phosphorylation in wildtype, *pib2Δ, pib2ΔNID,* or *pib2ΔCAD* cells during log phase growth (0), after 30 minutes of rapamycin treatment (rap), or complete nitrogen starvation (-N). **(E)** Phos-tag gel examining Ser33 phosphorylation in cells grown to mid-log phase and then starved for leucine (left gel) or starved for glutamine and treated with MSX (right gel). **(F)** Phos-tag gel examining Ser33 phosphorylation in cells grown to mid-log phase in media containing 0.5 g/L proline as the sole nitrogen source and then after addition of 0.5 g/L glutamine to the medium. **(G)** Co-immunoprecipitations showing an interaction between GFP-Pib2 and Ser33-HA (top panel), but not Gtr1-myc and Ser33-HA (bottom panel). Note we were not able to capture Pib2 from cells exposed to 2 hours of amino acid starvation. **(H)** Growth of wildtype, Ser33*^S7A^*, and Ser33*^S7D^* strains in synthetic medium missing serine and glycine. Cells were grown overnight in SD medium and then diluted into fresh medium missing serine and glycine at the start of the time-course. The lines and color matched shadows show the average and standard deviation from four replicates. Note that all strains are missing Ser3 to isolate the effect of Ser33.

**Figure 5 S1.**
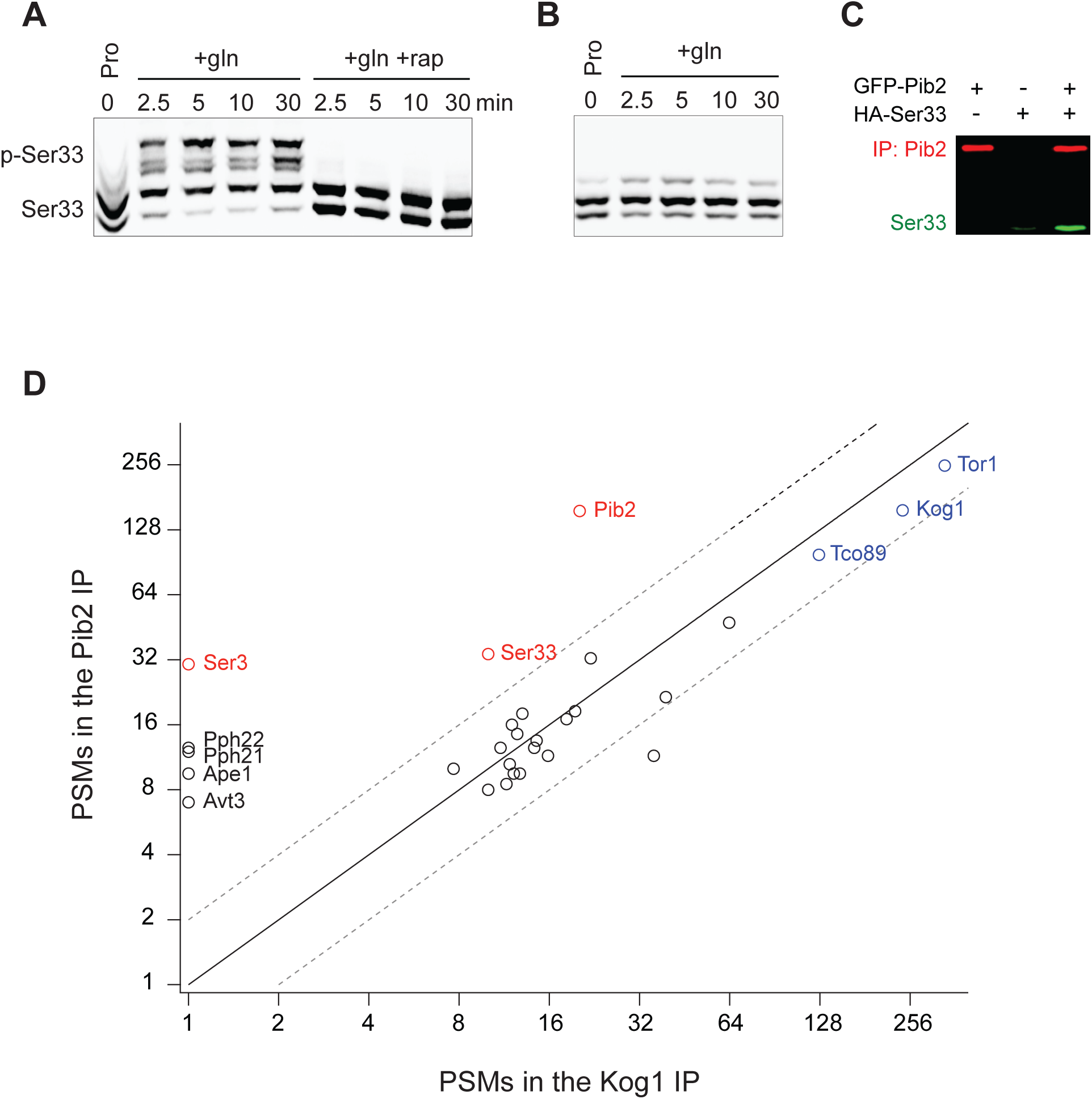
Rapid Ser33 phosphorylation by TORC1 in response to glutamine stimulation and evidence for a direct interaction with Pib2. **(A)** Phos-tag gel examining Ser33 phosphorylation in cells grown to mid-log phase in proline medium (time 0) and then after addition of 0.5 g/L glutamine (with or without rapamycin). **(B)** Phos-tag gel examining the phosphorylation of SER33^S7A^ grown to mid-log phase in proline medium (time 0) and then after addition of 0.5 g/L glutamine. **(C)** Co-immunoprecipitation showing an interaction between GFP-Pib2 and Ser33-HA during log phase growth (repeat of Fig. 5f). **(D)** Comparison between the amount of 30 proteins captured (as determined by the number of Peptide Spectral Maps) in a Pib2 immunopurification and a Kog1 immunopurification. Proteins are included on the graph if they were captured in both published Pib2 IPs ^73^ (many proteins are captured in a Kog1 IP, but not a Pib2 IP).

Next, we wanted to identify the stimuli that impact TORC1 signaling to Ser33. Using Phostag gel electrophoresis^72^, we found that rapamycin, glucose starvation, and complete nitrogen starvation (but not amino acid starvation) all lead to the rapid dephosphorylation of Ser33 (Fig. 5b). As expected, mutation of the seven TORC1 dependent phosphorylation sites listed above to alanine, or deletion of Pib2, also blocked the rapamycin dependent phosphorylation events detected on the gel (Figs. 5c, d).

The pattern of regulation seen for Ser33—with strong glucose, rapamycin and nitrogen, but limited amino acid, dependence—matched that seen previously for Sch9/Rps6^48,73^, leading to the question; what, if any, unique property (or properties) does TORC1-Pib2 dependent signaling to Ser33 have? To address this question, we first measured Ser33 phosphorylation during leucine and glutamine starvation. Here, we saw slow dephosphorylation in leucine starvation, and no change in glutamine starvation, just as with Rps6 (compare Fig. 5e and Fig. 1). We then wondered if TORC1-Pib2 signaling to Ser33 responds to the quality of the nitrogen source in the growth medium. To test this, we followed Ser33 phosphorylation in a prototrophic strain as it transitioned from growth in a poor-quality nitrogen source (proline medium) to a high-quality nitrogen source (glutamine medium)^74^. This experiment revealed that Ser33 phosphorylation is exquisitely sensitive to the quality of the amino acid/nitrogen source, with the TORC1 dependent sites going from unphosphorylated in proline, to >80% phosphorylated in glutamine (Figs. 5f, 5S1a, b). Other substrates we looked at (as detailed in Fig. 6) do not have such a strong response during the proline to glutamine shift, suggesting that the Pib2 dependence underlies this unique connection between TORC1 and Ser33.

**Figure 6:**
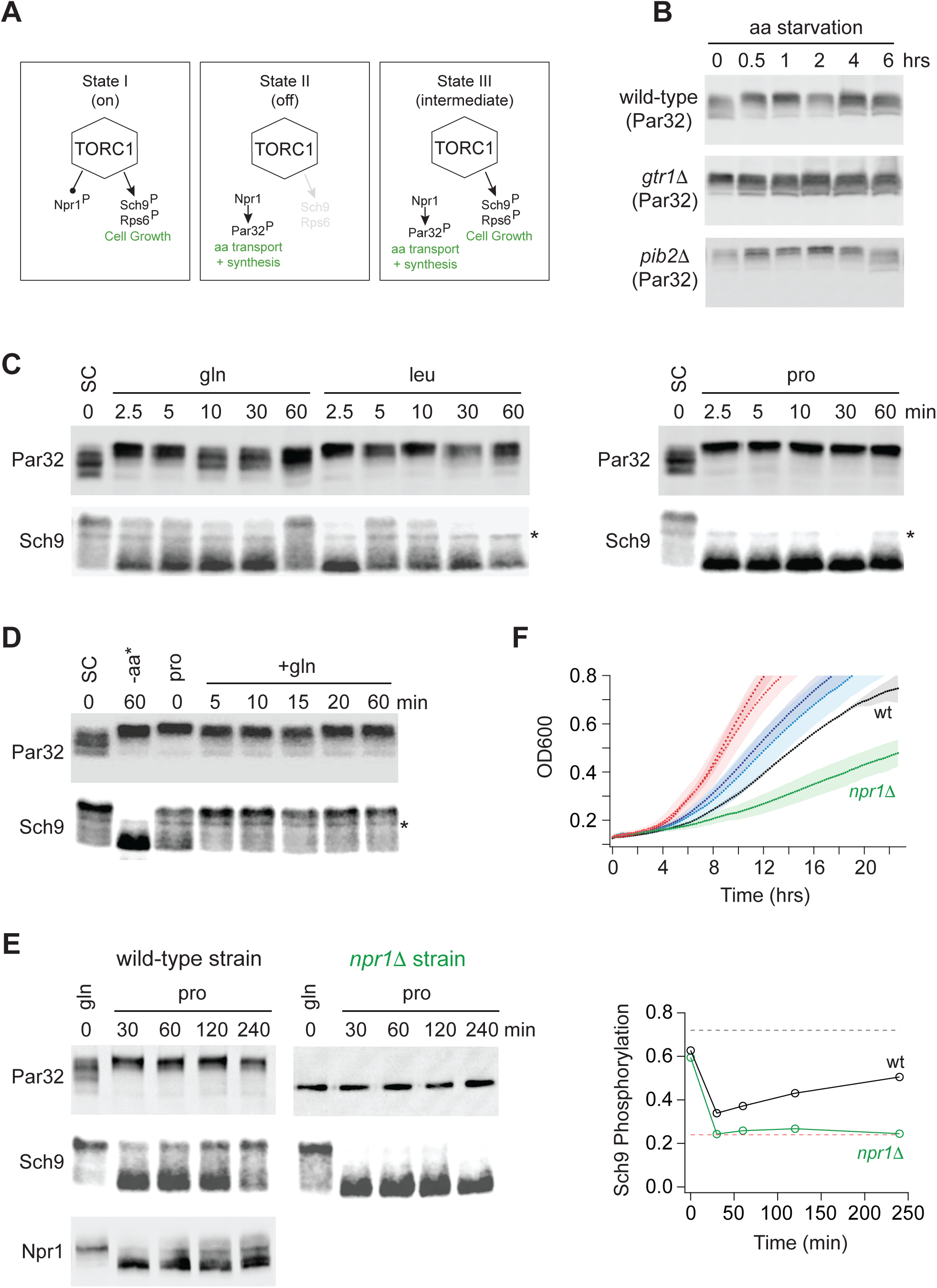
Growth in a poor nitrogen source pushes TORC1 into an intermediate signaling state. **(A)** Three state model of TORC1 signaling as described in the text. **(B)** Par32 phosphorylation measured by SDS-PAGE mobility shift during complete amino acid starvation in wildtype, *gtr1Δ*, and *pib2Δ* strains. **(C)** Par32 and Sch9 phosphorylation in cells grown to mid-log phase in synthetic complete media and then transferred to media containing either glutamine, leucine, or proline as the sole nitrogen source. The asterisk (*) highlights a non-specific band in the Western. **(D)** Par32 and Sch9 phosphorylation in cells grown to mid-log phase in synthetic complete medium (SC) and then exposed to complete amino acid starvation for 60 min (-aa) or grown in medium containing 0.5 g/L proline as the sole nitrogen source (pro) before 0.5 g/L glutamine was added to the culture. **(E)** Par32 and Sch9 phosphorylation in wild-type cells (left panel) and *npr1Δ* cells (right panel) grown to mid-log phase in medium containing 0.5 g/L glutamine as the sole nitrogen source (gln), and then transferred to media containing 0.5 g/L proline as the sole nitrogen source. The graphs on the right show the fraction of Sch9 phosphorylated at each time point, quantified by measuring the fraction of the Sch9 signal that runs above the fastest migrating band. The broken black and red lines show the fraction of Sch9 phosphorylated in SD medium, and after 1 hour of rapamycin treatment, respectively. **(F)** Growth of wildtype and *npr1Δ* cells in SC medium (wildtype dark red, *npr1Δ* light red), glutamine medium (wildtype dark blue, *npr1Δ* light blue), and proline medium (wildtype black, *npr1Δ* green). The lines and color matched shadows show the average and standard deviation from three replicates.

We also wanted to understand why the TORC1 dependent phosphorylation of Ser33 (and Ser3) requires Pib2. To address this question, we examined our recently published interactome data for Kog1 (the major regulatory subunit in TORC1) and Pib2^73^. This analysis revealed that Ser3 and Ser33 are two out of a total of six proteins that are captured at a significantly higher level in a Pib2 immunopurification than in a Kog1 immunopurification (out of >200 proteins enriched in the Kog1 IP), suggesting that Ser3 and Ser33 bind to Pib2 and not TORC1 (Fig. 5S1d). In line with this, we found that Ser33-HA is enriched in a Pib2 purification (2.5-fold) but not a Gtr1/2 purification (Fig. 5g).

Finally, to see if the TORC1 and Pib2 dependent phosphorylation of Ser33 is important for cell function, we followed the growth of phosphonull (S7A) and phosphomimic (S7D) versions of Ser33 in medium missing serine and glycine (in a Ser3 delete background). These experiments showed that the phosphonull (but not the phosphomimic) strain has a significant delay exiting quiescence, but then grows at the same rate as the wild-type strain once it is dividing rapidly (Fig. 5h). These data suggest (but do not prove) that the TORC1 and Pib2 dependent phosphorylation of Ser33 helps drive serine and glycine synthesis as cells transition into a rapid growth state, but TORC1 dependent phosphorylation is not required to maintain basal Ser33 activity during log phase growth.

### Multilevel signaling through TORC1

Our discovery that the deletion of Gtr1/2 or Pib2 leads to a change in TORC1 signaling through some substrates (particularly those involved in amino acid transport and metabolism), but not those involved in cell growth control, led us to hypothesize that the TORC1 pathway can take up at least three distinct signaling states (the first two of which are well known; Fig. 6a): (I) A fully active state to promote cell growth and inhibit the Npr1 dependent amino acid starvation response. (II) An inactive state to block cell growth and activate the Npr1 dependent amino acid starvation response. (III) A partially active state to simultaneously promote cell growth and activate the Npr1 dependent amino acid starvation response.

To test this model (i.e. see if State III exists in a wild-type strain), we needed reliable reporters of the TORC1 dependent cell growth and amino acid starvation responses. We therefore turned to two well established assays:

First, to follow the cell growth response, we monitored the phosphorylation of cleaved Sch9 using an SDS-PAGE mobility assay^48,54^. This assay follows the phosphorylation of several TORC1 target sites at the C-terminus of Sch9, known to play a key role in activating the kinase and protein synthesis. These C-terminal sites remain phosphorylated in the Gtr1/2 and Pib2 delete strains (Ser 723 and 726; Fig. 3 and right column Fig. 4).

Second, to follow the TORC1 dependent amino acid starvation response, we monitored the phosphorylation of full length Par32 using an SDS-PAGE mobility assay^48^. Par32 regulates amino acid transporter activity and is a substrate/reporter for the key TORC1 dependent amino acid starvation response regulator, Npr1^64^. Importantly, both Par32 and Npr1 are highly sensitive to the deletion of Gtr1/2 or Pib2 (left columns; Fig. 4 and 4S1). In the case of Par32, some sites are dephosphorylated in rapamycin and the mutant strains (Fig. 3 and left column, Fig. 4), while others (targeted by Npr1, a kinase that is repressed by TORC1) are hyperphosphorylated in rapamycin and the mutant strains (left column, Fig. 4S1). As a result, Par32 exists in a partially phosphorylated state in synthetic complete medium (SC) and then shifts to a hyperphosphorylated (slow migrating) state during amino acid starvation^48,64^ and in *gtr1*τι and *pib2*τι cells (Fig. 6b).

Once we established the Par32 and Sch9 reporter assays, we looked to see if transferring a prototrophic strain from medium that contains an excess of all 20 amino acids (SC), to medium containing a single high quality nitrogen source (glutamine) or a single poor quality nitrogen source (leucine or proline), pushes the cell into the predicted Sch9 on and Par32 on, intermediate signaling state (State III). This was not the case: In glutamine medium, the intermediate signaling state was populated for a short time (2.5 and 5 min; Fig. 6c), but Par32 was dephosphorylated again after 10 min (Fig. 6c). In leucine medium, we also saw transient population of the intermediate state, but here Sch9 was entirely dephosphorylated after 30 min (Fig. 6c). Finally, in proline medium, we saw a transition into the complete starvation state (State II), where Par32 is phosphorylated and Sch9 is dephosphorylated (Fig. 6c).

Next, we looked to see if the intermediate state is populated as cells transition from growth in a poor-quality nitrogen source (proline) to a high-quality nitrogen source (glutamine). This was the case, and to our surprise, TORC1 was already in the intermediate (Sch9 on, Par32 on) signaling state during steady state growth in proline medium (Fig. 6d). Thus, it appeared that the TORC1 pathway is driven into a complete starvation state (State II) when cells are first exposed to a poor nitrogen source (2.5-60 min, Fig. 6c) but then transitions into the intermediate signaling state (State III) as the cells adapt to the poor growth conditions (time 0, Fig. 6d).

To test this model further, we grew a prototrophic strain in glutamine medium and transferred it to proline medium, but this time followed Par32, Sch9, (and Npr1) phosphorylation for four hours. As predicted, Sch9 was dephosphorylated initially, but then reactivated (phosphorylated) over time (Fig 6e). In contrast, Par32 and Npr1 remained in an active or partially active form (highly phosphorylated and dephosphorylated, respectively) during the entire time-course (Fig. 6e).

We also carried out the same experiment in a *npr1*τι strain to test the prediction that the adaptation to a poor-quality nitrogen source is driven by the activation of Npr1 and Par32. This was the case, as Sch9 remained dephosphorylated during the entire time course in *npr1*τι cells (Fig. 6e).

The observation that cells growing at steady state in synthetic complete medium (SC), or glutamine medium, activate Sch9 and inhibit the Npr1-Par32 amino acid starvation response (Fig. 6c, e), while cells growing in proline medium activate both Sch9 and the Npr1-Par32 dependent amino acid starvation response (Fig. 6e), also led to another prediction. Wild-type and *npr1*τι cells should grow at the same rate in SC and glutamine medium, since the adaptive amino acid starvation response is off. However, in proline medium, where Npr1-Par32 signaling is active and presumably ensures that the cells can import/synthesize adequate amounts of nitrogen and each amino acid, the *npr1*τι cells should grow much slower than wild-type cells. This was also true (Fig. 6f).

### Tod6 is activated in the intermediate signaling state

The growth curves we collected in SC, glutamine and proline medium, highlight an important fact; yeast grow about four-times faster in SC medium than in proline medium (compare red and black lines, Fig. 6f). However, we see relatively little difference between the phosphorylation level of Sch9--the presumed master regulator of cell growth downstream of TORC1--in those two conditions (Fig. 6d). This observation led us to posit that one or more of the cell growth control factors is also part of the intermediate (State III) response and acts to slow cell growth in proline medium. The relevant cell growth regulators in yeast are: (i) Sfp1, a TORC1 dependent activator of ribosome and protein synthesis gene expression (and thus growth)^75,76^; (ii) Tod6, a TORC1 and Sch9 dependent repressor of ribosome and protein synthesis genes^77,78^; (iii) Stb3, another TORC1 and Sch9 dependent repressor of ribosome and protein synthesis genes^78,79^; (iv) Maf1, a TORC1 and Sch9 dependent repressor of RNA Polymerase III and tRNA degradation^49^. Therefore, to test our model, we searched our phosphoproteomics data to see if any of the aforementioned factors are sensitive to the deletion of Gtr1/2 or Pib2 (and thus likely activated/repressed in the intermediate state). We did not detect Sfp1 in our experiments, but found that Stb3 and Maf1 remain phosphorylated, while Tod6 is dephosphorylated, in the Gtr1/2 and Pib2 delete cells (Figs. 3 and 4).

Previous studies have shown that TORC1 inhibition triggers the movement of Stb3 and Tod6 into the nucleus where they act to recruit the Rpd3L deacetylase to, and thus repress, the ribosome protein and ribosome biogenesis genes^55^. Therefore, to test if Tod6 is part of the intermediate (State III) response, we measured the localization of Tod6-GFP and Stb3-GFP in a prototrophic strain carrying the nuclear marker Htb2-RFP and growing in SC medium, SC medium + rapamycin, or proline medium. As predicted, Tod6 moved into the nucleus in both proline medium and rapamycin, while Stb3 only moved into the nucleus in rapamycin (Fig. 7).

**Figure 7.**
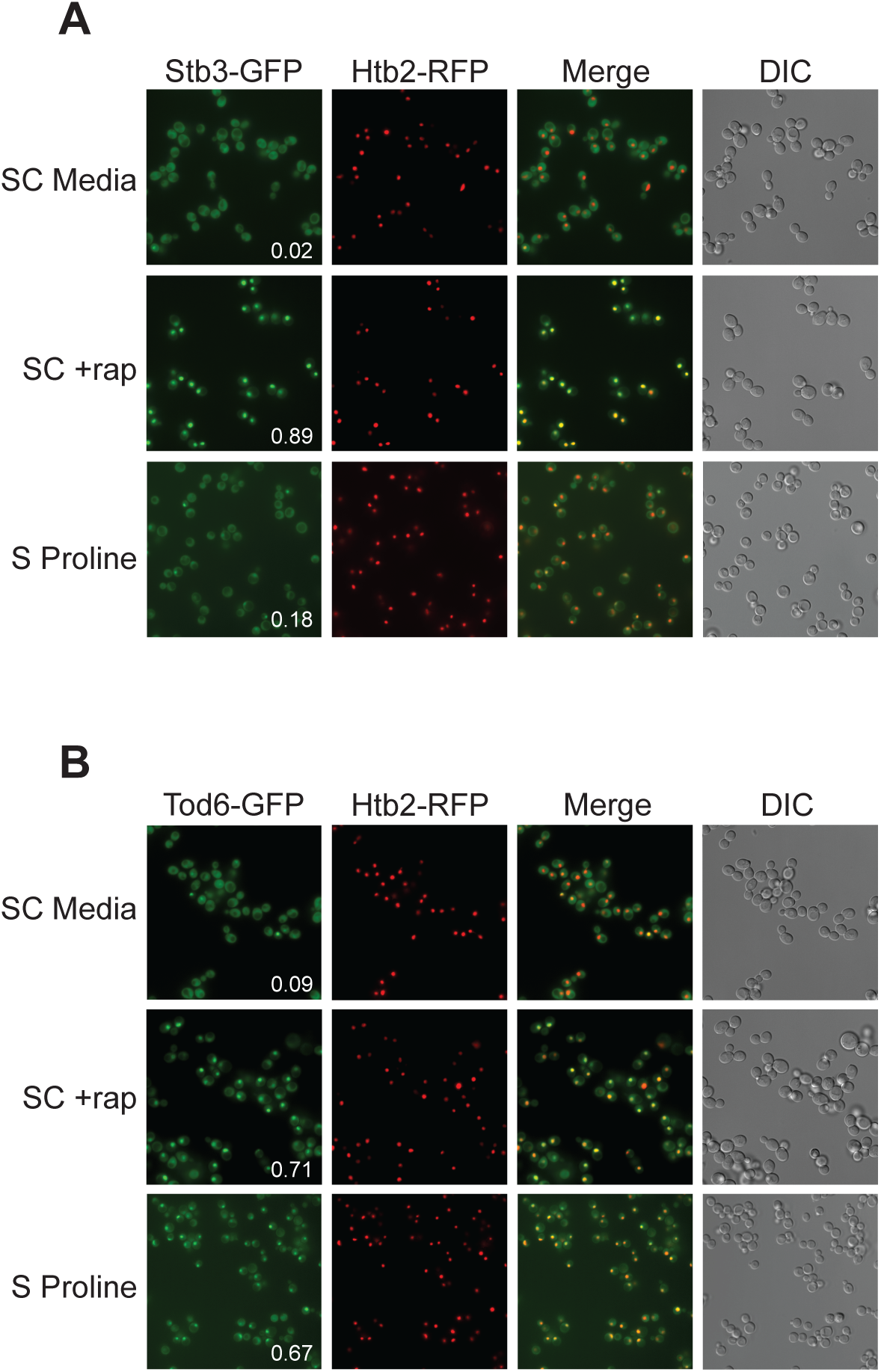
Tod6 moves to the nucleus during growth in a poor-quality nitrogen source. Localization of Stb3-GFP (top) and Tod6-GFP (bottom) during mid-log phase in SC medium, following exposure to 200 nM rapamycin for one hour, and during log phase in media containing 0.5 g/L proline as the sole nitrogen source (as labelled). The Tod6-GFP and Stb3-GFP strains are both prototrophic and carry the nuclear marker, Htb2-RFP, at its native locus. The numbers in the Stb3-GFP and Tod6-GFP panels show the fraction of cells with a strong nuclear GFP signal. In the SD and rapamycin control experiments these values are from a single experiment examining >200 cells. In the proline experiments the values are an average from three biological replicates, with >200 cells per replicate. In those three replicates, Stb3 was nuclear in 21% 17%, and 15% (18 ± 3%) of the cells, while Tod6 was nuclear in 70%, 70%, and 60% (67 ± 6%) of the cells.

Thus, Tod6 is dephosphorylated/activated in proline medium (State III), presumably to help slow the growth rate of the cell in the poor-quality nitrogen source.

### Gtr1/2 and Pib2 signaling during growth in a poor-quality nitrogen source

As a last step in our study, we wanted to see if, and how, changes in Gtr1/2 and Pib2 signaling drive TORC1 into the intermediate state. To address this question, we measured Par32 phosphorylation during the transition from growth in glutamine medium, to growth in proline medium, as we did earlier (Fig. 6e), but this time in a strain carrying mutations that lock Gtr1 and 2 into their active Gtr1-GTP and Gtr2-GDP bound forms (Gtr1^Q65L^ and Gtr2^S23L^, Gtr1/2^on^ for short^18^). In the Gtr1/2^on^ strain, Par32 was hyper-phosphorylated during the initial phase of the starvation response (when TORC1 is completely inactive; Fig. 6e), but then (erroneously) dephosphorylated over time (compare Fig. 8c and 6e), demonstrating that: (i) Gtr1/2 are normally inhibited during steady state growth in proline, and (ii) Gtr1/2 must remain inactive or partially inactive to keep TORC1 pathway in the intermediate (Par32 on) signaling state. We also measured the impact that locking Gtr1 and 2 in their inactive Gtr1-GDP and Gtr2-GTP bound forms (Gtr1/2^off^) has on Rps6 and Sch9 phosphorylation during steady state growth in proline medium and the transition to growth in glutamine medium. Locking Gtr1/2 off caused a moderate decrease in Sch9 (but not Rps6) phosphorylation (Fig. 8a, b), indicating that Gtr1/2 is partially (rather than fully) inactive during growth in proline medium.

**Figure 8.**
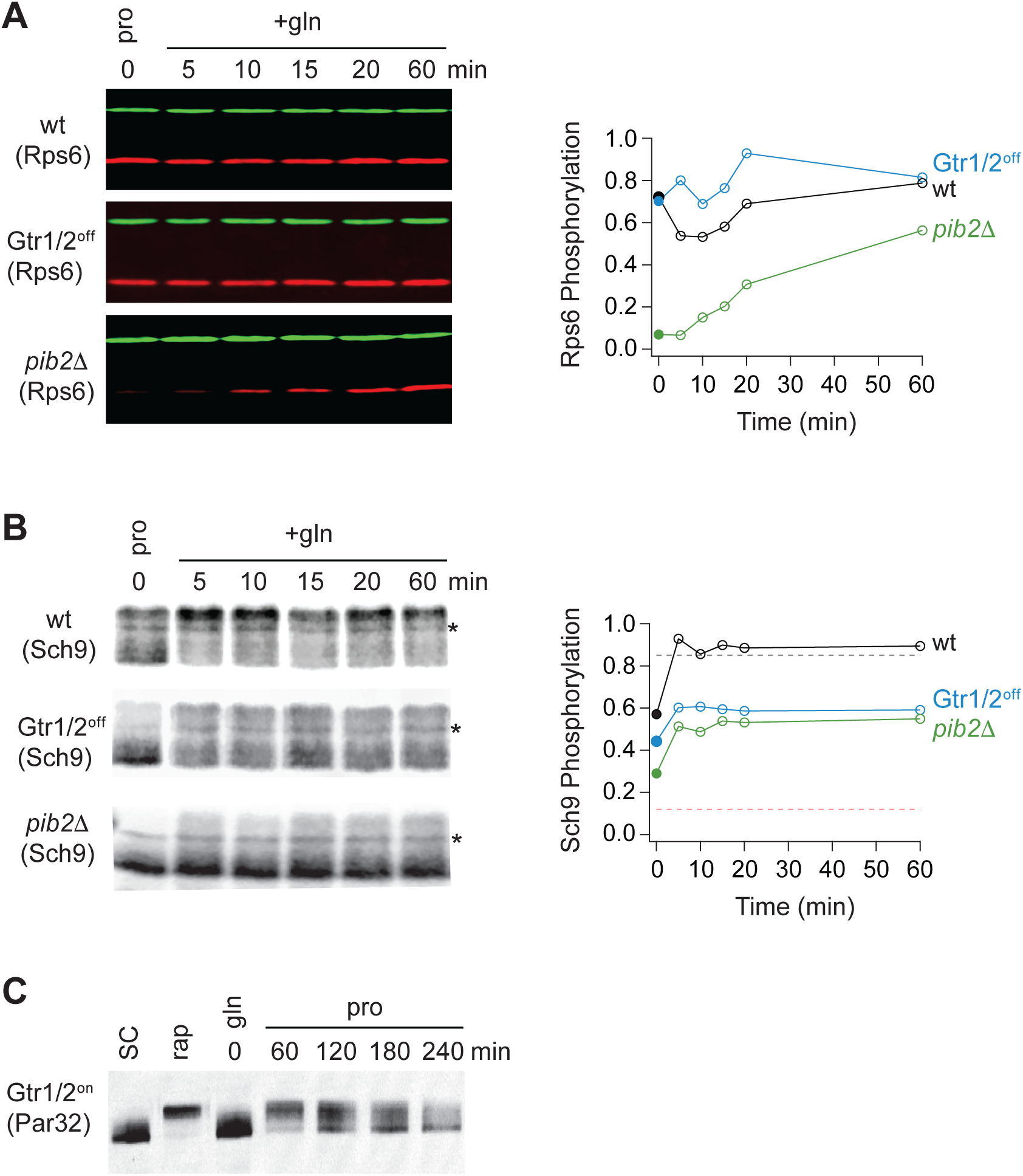
Partial Gtr1/2 inactivation drives TORC1 into the intermediate signaling state. **(A)** Rps6 and **(B)** Sch9 phosphorylation, measured in wild-type, GTR1/2^OFF^, and *pib2*11 strains, as cells transition from growth in proline medium to growth in glutamine medium. The data are quantified as described in Figs. 1 and 6e. Note, wild-type SC samples were included on each gel but were trimmed off the images presented for clarity. **(C)** Par32 phosphorylation as measured by SDS-PAGE mobility shift in GTR1/2^ON^ cells, either grown to mid-log phase in SC medium then treated with 200 nM rapamycin, or grown to mid-log phase in glutamine medium and then switched to proline medium (compare to Fig. 6e).

In a parallel set of experiments, we also examined the role that Pib2 plays in regulating TORC1 during growth in proline medium. This was more challenging since relatively little is known about the mechanisms underlying Pib2 signaling. Nevertheless, to assess the role of Pib2 in Npr1-Par32 activation we measured Par32 phosphorylation during the transition from growth in glutamine medium, to growth in proline medium, in a *pib2*τι*NID* strain (to lock Pib2 in an active form). Unexpectedly, Par32 was completely degraded in the absence of a NID domain in these conditions (data not shown). We have not seen the loss of Par32 in any natural condition, indicating that the NID domain of Pib2 is required for the normal function of the Npr1-Par32 branch of the TORC1 pathway, and that we cannot trap Pib2 in an active state. We then measured the impact that deleting Pib2 has on Rps6 and Sch9 phosphorylation during steady state growth in proline medium and the transition to growth in glutamine medium. These experiments showed that deleting Pib2 causes a large decrease in Rps6 and Sch9 phosphorylation at time zero (Fig. 8a, b) and thus that Pib2 is active, or mostly active, during growth in proline medium.

Thus, we conclude that Gtr1/2 is partially off, and Pib2 is on (or mostly on), during steady state growth in proline medium, and this pushes TORC1 into the intermediate/adaptive (Par32 on, Sch9 on) signaling state.

## DISCUSSION

In this report we show that the TORC1 pathway takes up at least three distinct signaling states (Fig. 9). In nutrient rich medium, TORC1 is fully active and phosphorylates (i) Sch9 and other proteins to drive cell growth, and (ii) Npr1 to suppress the amino acid starvation response (left panel, Fig. 9). In contrast, when cells are first transferred to medium containing a poor-quality nitrogen source like proline, TORC1 is inhibited (middle panel, Fig. 9). This blocks cell growth and triggers the activation of Npr1. Then, as cells adapt to the poor-quality nitrogen source (via Npr1), TORC1 is driven into an intermediate activity state where it phosphorylates Sch9 to re-initiate growth, but not Npr1, so that the cell continues to stimulate the increased amino acid transport and synthesis needed to support mass accumulation (right panel, Fig. 9). Tod6 is also dephosphorylated in the intermediate state, presumably to slow the growth rate of the cell so that it matches the maximum rate achievable in the poor-quality nitrogen source (right panel, Fig. 9).

**Figure 9.**
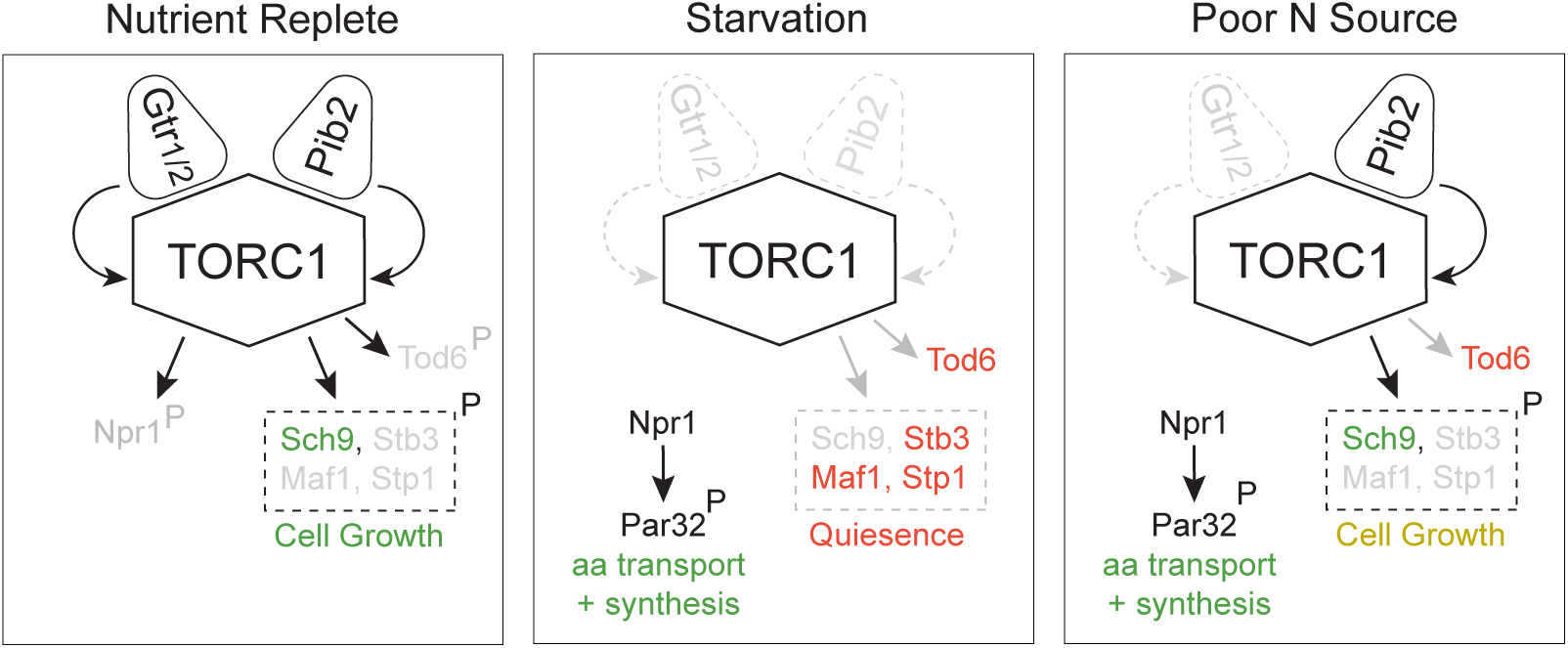
Three-state signaling through the TORC1 pathway. Schema describing the three signaling states of the TORC1 pathway, as described in the text. Grey arrows and names show inactive signaling events and proteins. Red names show active repressors; green names show active activators. Sch9, Stb3, Maf1 and Stp1 are phosphorylated in the same conditions and thus shown as a single functional unit (box with broken lines). Rapid cell growth is labeled green, while slow cell growth is labeled yellow.

The TORC1 pathway is driven into one of the three signaling states described above based on the combination of signals transmitted through Gtr1/2 and Pib2 (Fig. 9). In nutrient rich medium, Gtr1/2 and Pib2 are both on and push TORC1 into the fully active state. Then, when cells are first transferred to a poor-quality nitrogen source, Gtr1/2 and Pib2 turn off, or mostly off, triggering strong TORC1 inhibition. Finally, as the cells adapt to the new conditions, Pib2 is reactivated, so that TORC1 can phosphorylate most of the cell growth proteins but not Npr1, Tod6 and other proteins described in Fig. 4.

This three-state signaling system provides an elegant solution to a problem facing yeast as they transition between nutrient sources. Specifically, when yeast cells grow in nutrient-rich medium they focus on rapid growth via the import of key amino acids using highly selective transporters^80^. As a result, when amino acid levels fall, the cells do not know if there is a low-quality nitrogen source in the extracellular milieu, and do not have the capacity to import it even if it is there. Thus, the cells have to inactivate TORC1 and slow/halt their growth. However, once Npr1 is activated, the cells build the capacity to import, and grow using, a wide range of nitrogen containing compounds^66,80,81^. Then, if an adequate nitrogen source is available, Pib2 is turned back on so that the cell restarts growth while keeping the Npr1 dependent starvation response (and the import of alternate nitrogen sources) active.

Important implications of this three-state model include that: (i) yeast cells (likely including fungal pathogens) can be pushed to import alternative nitrogen sources, including toxic amino acid analogs and related compounds, by deletion or inhibition of Gtr1/2 or Pib2, and (ii) yeast cell growth on poor nitrogen sources can be halted by deletion of, or inhibition of, Pib2.

The data presented here also reveal another unique aspect of the Gtr1/2 and Pib2 control circuit, namely that the activity of Ser33 (and its homolog of Ser3) depends on signaling through TORC1-Pib2, and that this in turn appears to ensure that Ser33 is only phosphorylated in the presence of a high-quality nitrogen source. Our data suggests that this occurs due to direct binding of Ser33 to Pib2, leading to a model that is reminiscent of the RagC dependent binding and regulation of TFEB in human cells, where TFEB is regulated in response to amino acid signals (via RagC), but not the hormone signals transmitted to TORC1 through Rheb^82,83^. We hypothesize that the TORC1-Pib2 connection to Ser33 is setup this way to couple Ser33 activity to the level of glutamate/glutamine in the cell (rather than overall amino acid levels and other stress signals that regulate Gtr1/2) since the second step in serine synthesis involves the transfer of an amine group from glutamate to the Ser33 product, 3-phosphohydroxypyruvate^84^.

Beyond the implications this work has for understanding TORC1 signaling in yeast, our study also provides important insight into the design (and analysis) of other signaling pathways, including mTOR. Previous experiments examining TORC1 signaling in *S. cerevisiae*, including our own, focused on measuring TORC1 activity using one or two downstream reporters (usually pRps6 or pSch9). This led to the view that Gtr1/2 and Pib2 act as redundant activators of TORC1^36^. In reality, however, Gtr1/2 and Pib2 are semi-redundant activators of TORC1, and drive TORC1 into a fully active, partially active, or inactive state via on/on, off/on or off/off modes of Gtr1/2 and Pib2 signaling. Fully active TORC1 then phosphorylates all of its downstream targets, while partially active TORC1 phosphorylates a subset of its targets, to create a condition-dependent response. We argue that other stress and starvation signaling pathways likely work in a similar way, it is just that multi-level signaling has been overlooked since is difficult to detect using standard reporter assays.

## ACKNOWLEDGEMENTS

We thank Claudio De Virgilio for sharing GTR1 and 2 mutant plasmids, and Kyle Cunningham for sharing the GFP-Pib2 plasmid, used to make our mutant strains. We also thank Phil Gafken and Lisa Jones of the Fred Hutchinson Cancer Research Center’s Proteomics Facility for carrying out the Ser33 peptide mapping experiments. This work was supported by the National Institutes of Health (NIH) grants R01GM097329 and T32GM136536.

## MATERIALS AND METHODS

### Strain construction

All of the strains in this study were made in haploid (W303) *S. cerevisiae*, using standard methods^85,86^ and are listed in Table S3 (note there are two tabs). For strains grown in single nitrogen source media, we restored prototrophy using single copy plasmids containing the required complements (*LEU2, HIS3,* and/or *URA3*) and by integrating the *TRP1* gene into the genome^87^.

### Synthetic Medium

The experiments in Fig. 1-4 and most of Fig. 5 used standard auxotrophic (His-, Leu-) lab strains. These cells were grown in synthetic complete (SC) medium containing ammonium sulfate and all 20 amino acids, and then switched to the same medium missing one or more amino acid, as indicated. The experiments in Fig. 5f and 6-8 used prototrophic yeast strains, and cells were grown and studied in synthetic medium without ammonium sulfate, supplemented with a single nitrogen source (e.g, Leu, Gln, or Pro) unless noted (SC panels, Fig. 7).

### Rps6 phosphorylation assay

Cultures were grown in conical flasks, shaking at 200 rpm and 30 °C until mid-log phase (OD_600_ 0.4-0.7). At that point, a 47 mL sample was collected, mixed with 3 mL of 100% trichloroacetic acid (TCA), and held on ice for at least 30 mins (but no more than 6 hours). The remaining culture was then collected by filtration, and transferred to SC -amino acids, SC -leu, or SC -glutamine medium after two washes with 100 ml of the same medium, and additional samples collected as described above. Cells exposed to glutamine starvation were also treated with a freshly made 2 mM methionine sulfoximine (MSX, Sigma Aldrich 15985-39-4). The TCA precipitated samples were then centrifuged at 4000 rpm for 5 min at 4°C, washed twice with 4°C water, twice with acetone, and disrupted by sonication (2X) at 15% amplitude for 5 s before centrifugation at 12,000 rpm for 30 s. The cell pellets were then dried in a speedvac for 10 min at room temperature, and frozen until required at −80°C.

Protein extraction was performed by bead beating (6 × 1 min, full speed) in Urea buffer (6 M Urea, 50 mM Tris–HCl pH 7.5, 5 mM Ethylenediaminetetraacetic acid (EDTA), 1 mM Phenylmethylsulfonyl fluoride (PMSF), 5 mM NaF, 5 mM NaN_3_, 5 mM NaH_2_PO_4_, 5 mM *p*-nitrophenylphosphate, 5 mM β-glycerophosphate, and 1% SDS) supplemented with complete protease and phosphatase inhibitor tablets (Roche, Indianapolis, IN; 04693159001 and 04906845001). The lysate was then harvested by centrifugation for 5 min at 3000 rpm, resuspended into a homogenous slurry, and heated at 65 °C for 10 min. The soluble proteins were then separated from insoluble cell debris by centrifugation at 12,000 rpm for 10 min, and the lysate stored at −80 °C until required.

For protein phosphorylation analysis, the protein extracts were run on a 12% acrylamide gel and transferred to a nitrocellulose membrane. Western blotting was then carried out using anti-pRPS6 antibody (Cell Signaling, 4858) at a 1/2500 dilution, and anti-PGK1 antibody (Invitrogen, 459250) at a 1/10,000 dilution, and anti-mouse and anti-rabbit secondaries, labeled with a IRDye 680RD (LiCor 926-68071) and IRDye 800CW (LiCor 926-32210) both at a 1/10,000 dilution, and the blots scanned using a LiCor Odyssey Scanner (LiCor, Lincoln, NE). Band intensities were quantified using the LiCor Image Studio Software.

### SYTOX Cell Death Assay

Cells were inoculated into a 10 mL starter culture, and then re-innoculated into SC medium to ensure at least 12 hours of logarithmic growth. Once the cells had reached an OD_600_ between 0.4 and 0.7 they were collected by filtration, washed with 25 mL of SC medium missing glutamine and supplemented with 2mM methionine sulfoximine (SC-gln), and then resuspended in 25 mL of the same medium for 6 hours. The cells were then moved into 8-well microslides (Ibidi, 80826) pretreated with Concanavalin A (Fischer Scientific ICN15071001) and stained with SYTOX Green Fluorescent Dye (Inivtrogen S7020). Specifically, after the cells settled in the well, they were washed with 50 mM Sodium Citrate, and then treated with 50 mM Sodium Citrate containing 1 mM SYTOX Green Dye for 30 minutes (in the dark). The cells were then washed two additional times with 50 mM Sodium Citrate, resuspended in the same buffer, and fluorescence and DIC images acquired using a Nikon Eclipse Ti-E microscope equipped with a 100X objective and a Photometrics Prime 95B camera (excitation at 488 nm, emission at 515 nm, 1 s exposure).

### Sch9, Par32, and Npr1 mobility shift experiments

Cells examined in the band shift (gel mobility) experiments were grown to OD_600_ 0.4-0.7 in SC medium, or SC medium missing amino acids but containing proline, glutamine, or leucine, as indicated. Samples were then collected by TCA precipitation and lysed in Urea buffer as described above for Rps6. Sch9-3xHA samples were also subjected to cleavage by 2-nitro-5-thiocyanatobenzoic acid (NTCB) in N-Cyclohexyl-2-aminoethanesulfonic acid (CHES, pH 10.5) overnight at 30 °C (1 mM NTCB and 100 mM CHES) as described previously ^48,88^.

Par32-13xMyc and Npr1-3XFLAG samples (25 and 100 μg of total protein, respectively) were run on a 7.5% acrylamide gel for 3 hours at 80 V and transferred to a nitrocellulose membrane. Western blotting was then carried out using anti-Myc (Thermo Scientific MA12316) or anti-FLAG (Sigma aldrich F1804-1MG) antibodies at a 1/1000 dilution, and an anti-mouse secondary labeled with a IRDye 800CW (LiCor 926-32210), at a 1/10,000 dilution, and the blots scanned using a LiCor Odyssey Scanner (LiCor, Lincoln, NE).

Sch9-3XHA samples were (10 μg of total protein, respectively) run on a 12% acrylamide gel for 3 hours at 80 V and transferred to a nitrocellulose membrane. Western blotting was then carried out using an anti-HA antibody (Sigma Aldrich, 11583816001) at a 1/1000 dilution, and an anti-mouse secondary, labeled with a IRDye 800CW (LiCor 926-32210), at a 1/10,000 dilution, and the blots scanned using a LiCor Odyssey Scanner (LiCor, Lincoln, NE).

### Ser33 PhosTag bandshift experiments

Ser33-3xFLAG cells were grown and harvested as described above for the Rps6 experiments. 25 μg of Ser33-3xFLAG protein extract (per lane), was then loaded on an 8% Zn^2+^/PhosTag BisTris gel at 60 V for 3 hours and 45 minutes in MOPS running buffer made following manufacturer’s instructions (FUJIFILM, Wako AAL-107). To prevent the 5 mM EDTA in our Urea buffer from disrupting band migration, we added 2uM ZnNO_3_ to our loading buffer (EDTA-free) prior to mixing it with our samples. After electrophoresis, the gels were washed with transfer buffer containing 5 mM EDTA for 10 minutes and then again with standard transfer buffer (no EDTA) for 10 minutes. Western blotting then proceeded as described above.

### Phosphoproteomics

The cells examined using phosphoproteomics were collected using TCA precipitation, as described above. All remaining steps used MS-grade reagents (including water):

First, the cell pellets were resuspended in 400 μL of MS-urea buffer (8M urea, 100mM ammonium bicarbonate (ABC), 5mM EDTA). Proteins were then extracted by bead beating (as described above) and eluted into wide-mouth tubes (without a 65 °C denaturation step) and the final protein concentration in each sample measured using a BCA assay.

200 μg of total protein was taken from each sample and diluted to 1 μg/μL in a 2.0 mL Low-bind tube (Thermo Scientific 88379) using the 8 M urea buffer above. The samples were alkylated and reduced by treatment with 5 mM Tris(2-carboxyethyl)phosphine hydrochloride (TCEP) and 5 mM iodoacetamide at room temperature for 30 min. The reduced and alkylated samples were then diluted by adding 70 μL of 50 mM ABC (so that the final urea concentration was 5.5 M) and 20 ng/μL of LysC (New England Biolabs, P8109S) to each sample, and digested at 37 °C, sharking at 700 rpm, for 3 hours. The samples were then diluted again with 1.3 mL 50 mM ABC to bring the urea concentration to 1 M, and treated with 2 μg trypsin (Promega, v511c), shaking overnight at 700 rpm, and 37 °C.

The next morning trypsinization was quenched by adding TFA to a final concentration of 1% v/v, and the samples clarified by centrifugation at 15,000 rpm for 5 minutes. The peptide mix was then desalted using Sep Pak Plus C18 cartridges (Waters: WAT020515) on a vacuum manifold. First, cartridges were equilibrated by flushing with 5 mL of solution B (65% acetonitrile (MeCN), 0.1% trifluoroacetic acid (TFA)) and then 10 mL solution A (2% MeCN, 0.1% TFA). The peptide samples, now around 2 mL in volume, were then diluted with 8 mL of solution A and slowly run through the cartridges. The columns were then washed with 10 mL of solution A and then peptides eluted twice using 600 μL of solution B and collected in a Lo-Bind tube. The peptides were then dried in a speed vac at room temperature and stored at -80 °C.

Phosphopeptides were enriched using magnetic Ti(IV)-IMAC beads (MagReSyn MR-TIM005) following the manufacturer’s instructions. Specifically, 40 μL beads were equilibrated using three washes with loading buffer (0.1 M glycolic acid in 80% MeCN, 5% TFA). Dried peptide samples were then resuspended in 200 μL loading buffer and incubated with the beads for 20 minutes, shaking at 600 rpm and room temperature. The beads were then washed with 200 μL of loading solution, 100 μL wash solution 1 (80% ACN, 1% TFA), and 100 μL wash solution 2 (10% ACN, 0.2% TFA) for 2 minutes each, with 600 rpm agitation. The phosphopeptides were then eluted twice using 135 μL of 1% NH_4_OH into 90 μl of 10% formic acid (FA), leading to a final volume of 360 μL.

The purified phosphopeptides were then desalted on micro spin columns (Nest Group: SEM SS18V). Each step used centrifugation at 1,500 g for 1 minute. First, columns were conditioned with 400 μl 90% MeCN, 0.1% FA and then equilibrated with 350 μL 5% MeCN, 0.1% FA. The samples were then loaded onto the columns, washed with 350 μL 5% MeCN, 0.1% FA, eluted in 200 μL 50% MeCN, 0.1% FA, dried using a speedvac at room temperature, and stored at -80 °C.

### Mass Spectrometry

Samples were resuspended in 10.5 μL 0.1% FA, and 1.5 μL of the suspension injected for HPLCESI-MS/MS analysis. Data acquisition was performed in positive ion mode on a ThermoScientific Orbitrap Fusion Lumos tribrid mass spectrometer fitted with an EASY-SpraySource (Thermo Scientific, San Jose, CA). NanoLC was performed using a ThermoScientific UltiMate 3000 RSLCnano System with an EASY Spray C18 LC column (ThermoScientific, 50 cm x 75 μm inner diameter, packed with PepMap RSLC C18 material, 2 μm, cat. # ES803): loading phase for 15 min at 0.300 μl/min; linear gradient of 1–34% Buffer B in 119 min at 0.220 μl/min, followed by a step to 95% Buffer B over 4 min at 0.220 μL/min, hold 5 min at 0.250 μl/min, and then a step to 1% Buffer B over 5 min at 0.250μl/min and a final hold for 10 in (total run 159 min); Buffer A = 0.1% FA; Buffer B = 0.1%FA in 80% ACN. Spectra were collected using XCalibur, version 2.3 (ThermoFisherScientific). Precursor scans were acquired in the Orbitrap at 120,000 resolution on a mass range from 375 to 1575 Th. Precursors were isolated with an isolation width of 1.6 Th and subjected to higher energy collisional dissociation. MS/MS scans were acquired in the ion trap on the m/z range of 120 to 2000 Th with a fill time of 35 ms.

### Phosphoproteomic data analysis

Progenesis QI for proteomics software (Version 2.4, Nonlinear Dynamics Ltd., Newcastle upon Tyne, UK) was used to perform ion-intensity based label-free quantification as described previously. In brief, in an automated format, .raw files were imported and converted into two-dimensional maps (y-axis = time, x-axis =m/z) followed by selection of a reference run for alignment purposes. An aggregate data set containing all peak information from all of the samples in a given experiment was created from the aligned runs, which was then further narrowed down by selecting only +2, +3, and +4 charged ions for further analysis. A peak list of fragment ion spectra was exported in Mascot generic file (.mgf) format and searched against a UniProt *S. cerevisiae* S288c database (6728 entries) using Mascot (Matrix Science, London, UK; version 2.6). The search variables that were used were: 10 ppm mass tolerance for precursor ion masses and 0.5 Da for product ion masses; digestion with trypsin; a maximum of two missed tryptic cleavages; variable modifications of oxidation of methionine and phosphorylation of serine, threonine, and tyrosine; 13C=1. The resulting Mascot .xml file was then imported into Progenesis, allowing for peptide/protein assignment, while peptides with a Mascot Ion Score of <25 were not considered for further analysis.

Peptide ion data were exported as a .csv file. Positions of phosphorylation sites within the protein were obtained by mapping the peptide sequence to the protein sequence in the .fasta file used for the database search. Duplicate entries of the same peptide ion mapping to more than one protein were collapsed to one entry. Normalized intensities of phosphorylated peptides mapping to the same phosphosite were summed together.

### Crosslinking and coimmunoprecipitation

Strains carrying GFP-Pib2 and 3xHA-Ser33, or Gtr1-13xMyc and 3xHA-Ser33, were grown in 500 mL of synthetic complete media to log phase as described above. 250 mL of each sample was collected by filtration and snap frozen in liquid nitrogen. The remaining samples were then captured by filtration, washed with 250 mL of SC medium missing amino acids (-aa), transferred to 250 mL of SC -aa media for 2 hours, and collected by filtration and snap freezing. To begin protein extraction, frozen cell pellets were washed with 5 mL of 4 °C Immunoprecipitation Lysis Buffer (IPLB; 20 mM 4-(2-hydroxyethyl)-1-piperazineethanesulfonic acid (HEPES), pH 7.5, 150 mM potassium acetate, 2 mM magnesium acetate, 1 mM ethylene glycol bis(2-aminoethyl)tetraacetic acid (EGTA), and 0.6 M sorbitol) (Murley et al., 2017) and then resuspended in 1 mL of IPLB supplemented with complete protease and phosphatase inhibitors (IPLB^++^). Samples resuspended in IPLB^++^ were split between two 2 mL screw-cap tubes, sheared by bead beating, and eluted into 1.5 mL wide-mouth tubes as described above. Lysates were homogenized by gentle vortexing and combined into a fresh 2 mL tube. Crosslinking was performed by treating the lysates with 0.25 μM of dithiobis(succinimidyl propionate) (DSP) at 4 °C for 30 minutes with gentle rotation. The reaction was then quenched by adding 100 mM Tris, pH 7.5, and the sample held on ice for 30 minutes. Cell membranes were then solubilized by adding digitonin to a concentration of 1% with gentle rotation for 30 minutes and clarified by centrifugation at 12,000 rpm at 4°C for 10 minutes, and the supernatants transferred to a fresh 2.0 mL tube.

To co-purify GFP-Pib2 or Gtr1-13xMyc and any interacting Ser33, 50 μL of μMACS anti-GFP or anti-c-Myc (Miltenyi Biotec, 130-091-125 and 130-091-123) was added to clarified extract and incubated at 4 °C while rotating for 2 hours. μMACS columns were equilibrated by adding 200uL’s of IPLB^++^ containing 1% digitonin. Samples were then added to each column (on a magnet) and allowed to pass through by gravity. The beads were then washed three times with 200μL of IPLB^++^ containing 0.1% digitonin, and then twice with 500 μL IPLB (without digitonin). The protein was then eluted, first by incubation for 5 minutes with 20 μL elution buffer (supplied with μMACS kit) pre-heated to 95°C, and then by adding 2×40 μL of the same buffer.

### Immunoprecipitation and MS-phosphomapping of Ser33

Immunoprecipitation of Ser33-3xFLAG was performed using the protocol described above, but without the DSP crosslinking, Tris quenching, and digitonin membrane-permeabilization steps. Instead, IPLB buffer was supplemented with 0.25% TWEEN, and 3xFLAG-tagged Ser33 was purified using anti-FLAG conjugated antibodies (130-101-591). Immunopurified samples were then separated by SDS-PAGE gel, and gel slices around bands corresponding to the correct molecular weight of Ser33 were excised and sent for phosphoproteomic analysis by mass spectrometry.

Gel slices were washed for 15 min each with water, 50/50 acetonitrile/water, acetonitrile, 100 mM ammonium bicarbonate, followed by 50/50 acetonitrile/100 mM ammonium bicarbonate. The solution was then removed and the gel slices dried by vacuum centrifugation. Next, the dried gel slices were reduced by covering them with 10 mM dithiothreitol in 100 mM ammonium bicarbonate and heating them at 56°C for 45 min; alkylated by covering them with a solution of 55 mM iodoacetamide in 100 mM ammonium bicarbonate and incubating in the dark at ambient temperature for 30min, and washed with 100 mM ammonium bicarbonate for 10 min and 50 mM ammonium biocarbonate + 50% acetonitrile for 10 min. The gel slices were then dried again and treated with an ice-cold solution of 12.5 ng/μL trypsin (Promega, Madison, WI) in 100 mM ammonium bicarbonate. After 45 min, the trypsin solution was removed, discarded, and a volume of 50 mM ammonium bicarbonate was added to cover the gel slices and they were incubated overnight at 37°C with mixing on a shaker. Samples were then spun down in a microfuge and the supernatant collected. The same Gel slices were then incubated in 0.1% trifluoroacetic acid (TFA) and acetonitrile, centrifuged, and the supernatant collected. At this point, the digestion supernatant and the extraction supernatant were pooled, split into two tubes, and concentrated using vacuum centrifugation. One tube was further digested with thermolysin (Promega, Madison, WI) by resuspending the tryptically digested peptides with a solution containing 50 mM Tris-HCl pH 8 and 0.5 mM calcium chloride and adding 1 μg of thermolysin. Digestion was carried out at 75°C with mixing for 5 hr. The thermolysin was quenched by adding TFA to a 0.5% final concentration. All samples were desalted using ZipTip C_18_ (Millipore, Billerica, MA) and eluted with 70% acetonitrile/0.1% TFA. The desalted material was concentrated to dryness in a speed vac. The proteolytically-digested samples were brought up in 20 μL of 2% acetonitrile in 0.1% formic acid and 18 μL and then analyzed by LC/ESI MS/MS with a Thermo Scientific Easy-nLC II (Thermo Scientific, Waltham, MA) coupled to a Orbitrap Elite ETD (Thermo Scientific, Waltham, MA) mass spectrometer using a trap-column configuration as described in ^89^. In-line de-salting was accomplished using a reversed-phase trap column (100 μm × 20 mm) packed with Magic C_18_AQ (5-μm, 200Å resin; Michrom Bioresources, Auburn, CA) followed by peptide separations on a reversed-phase column (75 μm × 250 mm) packed with Magic C_18_AQ (5-μm, 100Å resin; Michrom Bioresources, Auburn, CA) directly mounted on the electrospray ion source. A 45-min gradient from 2% to 35% acetonitrile in 0.1% formic acid at a flow rate of 400 nL/min was used for chromatographic separations. A spray voltage of 2500V was applied to the electrospray tip and the Orbitrap Elite instrument was operated in the data-dependent mode, switching automatically between MS survey scans in the Orbitrap (AGC target value 1,000,000, resolution 120,000, and injection time 250 milliseconds) with collision induced dissociation MS/MS spectra acquisition in the linear ion trap (AGC target value of 10,000 and injection time 100 milliseconds), higher-energy collision induced dissociation (HCD) MS/MS spectra acquisition in the Orbitrap (AGC target value of 50,000, 15,000 resolution and injection time 250 milliseconds) and electron transfer dissociation (ETD) MS/MS spectra acquisition in the Orbitrap (AGC target value of 50,000, 15,000 resolution and injection time 250 milliseconds). The three most intense precursor ions from the Fourier-transform (FT) full scan were consecutively selected for fragmentation in the linear ion trap by CID with a normalized collision energy of 35%, fragmentation in the HCD cell with normalized collision energy of 35%, and fragmentation by ETD with 100 ms activation time. Selected ions were dynamically excluded for 30 seconds.

Data analysis was performed using Proteome Discoverer 1.4 (Thermo Scientific, San Jose, CA). The data were searched against the Saccharomyces Genome Database (downloaded 02/03/2011; www.yeastgenome.org) that was appended with protein sequences from the common Repository of Adventitious Proteins or cRAP (www.thegpm.org/crap/). Two searches were performed corresponding to the proteolytic enzymes trypsin, and trypsin/thermolysin (no enzyme selected). The Maximum missed cleavages was set to 2. The precursor ion tolerance was set to 10 ppm and the fragment ion tolerance was set to 0.8 Da. Variable modifications included oxidation on methionine (+ 15.995 Da), carbamidomethyl (+ 57.021 Da) on cysteine, and phosphorylation on serine, threonine, and tyrosine (+ 79.996 Da). Sequest HT was used for database searching. PhosphoRS 3.1 ^90^ was used for assigning phosphosite localization probabilities. All search results were run through PSM Validator for false discovery rate evaluation of the identified peptides.

### Fluorescence microscopy

For microscopy in SC + proline, cells were grown in 25 mL over three days, refreshing the media by removing half and re-adding equivalent volume each day, until they reached the logarithmic growth phase. Cells grown in SC were inoculated into a 10 mL starter culture, and then diluted again into a 25 mL overnight culture. In both cases, cells were grown to log phase (OD_600_ 0.4-0.6) and then pipetted onto 8-well microslides (Ibidi, 80826) that had been pretreated with Concanavalin A. Cells were then washed with matching media (SD + proline or SC) and then imaged using a Nikon Eclipse Ti-E microscope equipped with a 100X objective and a Photometrics Prime 95B camera. GFP images were acquired with an excitation of 488 nm and an emission of 515 nm using 1 second exposure. RFP images were acquired with an excitation of 561 nm and an emission of 632 nm using a 500 ms exposure. DIC images were captured using a 60 ms exposure. To examine the impact of complete TORC1 inhibition, cells growing in SC medium were treated 200 mM rapamycin for 60 min (within the well) and then imaged again with the same settings. Images were analyzed and quantified in ImageJ ^91^.

### Proliferation assays

3 biological replicates of each strain was grown overnight and aliquoted to tubes containing 1 OD_600_ unit. Samples were then washed with 1 mL filtered water, and then resuspended in 1 mL filtered water. 100 μL of culture were diluted into 900 μL of each medium, bringing the starting concentration to an OD_600_ of 0.1. For each biological replicate, 3 technical replicates were plated across a 96-well plate (Corning 3370). Plates were covered with Breathe-Easy membranes (Z380059 Sigma-Aldrich) and growth was measured on a TECAN Infinite M Nano plate reader. Growth settings were orbital shaking with a 5 mm amplitude for 8 minutes followed by a linear shake with a 5 mm amplitude for 2 minutes at 30°C. OD_600_ readings were taken every 10 minutes for 23 hours.

## REFERENCES

1. Loewith, R., and Hall, M.N. (2011). Target of Rapamycin (TOR) in Nutrient Signaling and Growth Control. Genetics 189, 1177–1201. 10.1534/geneLcs.111.133363.

2. Liu, G.Y., and Sabatini, D.M. (2020). mTOR at the nexus of nutrition, growth, ageing and disease. Nature Reviews Molecular Cell Biology 21, 183–203. 10.1038/s41580-019-0199-y.

3. González, A., and Hall, M.N. (2017). Nutrient sensing and TOR signaling in yeast and mammals. The EMBO Journal 36, 397–408. 10.15252/embj.201696010.

4. Robitaille, A.M., Christen, S., Shimobayashi, M., Cornu, M., Fava, L.L., Moes, S., Prescianotto-Baschong, C., Sauer, U., Jenoe, P., and Hall, M.N. (2013). Quantitative Phosphoproteomics Reveal mTORC1 Activates de Novo Pyrimidine Synthesis. Science 339, 1320–1323. doi:10.1126/science.1228771.

5. Peterson, T.R., Sengupta, S.S., Harris, T.E., Carmack, A.E., Kang, S.A., Balderas, E., Guertin, D.A., Madden, K.L., Carpenter, A.E., Finck, B.N., and Sabatini, D.M. (2011). mTOR complex 1 regulates lipin 1 localization to control the SREBP pathway. Cell 146, 408–420. 10.1016/j.cell.2011.06.034.

6. Huber, A., Bodenmiller, B., Uotila, A., Stahl, M., Wanka, S., Gerrits, B., Aebersold, R., and Loewith, R. (2009). Characterization of the rapamycin-sensitive phosphoproteome reveals that Sch9 is a central coordinator of protein synthesis. Genes Dev 23, 1929–1943. 10.1101/gad.532109.

7. Hsu, P.P., Kang, S.A., Rameseder, J., Zhang, Y., Ofna, K.A., Lim, D., Peterson, T.R., Choi, Y., Gray, N.S., Yaffe, M.B., et al. (2011). The mTOR-Regulated Phosphoproteome Reveals a Mechanism of mTORC1-Mediated Inhibition of Growth Factor Signaling. Science 332, 1317–1322. doi:10.1126/science.1199498.

8. Ben-Sahra, I., and Manning, B.D. (2017). mTORC1 signaling and the metabolic control of cell growth. Current Opinion in Cell Biology 45, 72–82. 10.1016/j.ceb.2017.02.012.

9. Ben-Sahra, I., Hoxhaj, G., Ricoult, S.J.H., Asara, J.M., and Manning, B.D. (2016). mTORC1 induces purine synthesis through control of the mitochondrial tetrahydrofolate cycle. Science 351, 728–733. doi:10.1126/science.aad0489.

10. Kim, J., Kundu, M., Viollet, B., and Guan, K.-L. (2011). AMPK and mTOR regulate autophagy through direct phosphorylation of Ulk1. Nature Cell Biology 13, 132–141. 10.1038/ncb2152.

11. Kamada, Y., Funakoshi, T., Shintani, T., Nagano, K., Ohsumi, M., and Ohsumi, Y. (2000). Tor-Mediated Induction of Autophagy via an Apg1 Protein Kinase Complex. Journal of Cell Biology 150, 1507–1513. 10.1083/jcb.150.6.1507.

12. Düvel, K., Yecies, J.L., Menon, S., Raman, P., Lipovsky, A.I., Souza, A.L., Triantafellow, E., Ma, Q., Gorski, R., Cleaver, S., et al. (2010). Activation of a Metabolic Gene Regulatory Network Downstream of mTOR Complex 1. Molecular Cell 39, 171–183. 10.1016/j.molcel.2010.06.022.

13. Sancak, Y., Peterson, T.R., Shaul, Y.D., Lindquist, R.A., Thoreen, C.C., Bar-Peled, L., and Sabatini, D.M. (2008). The Rag GTPases Bind Raptor and Mediate Amino Acid Signaling to mTORC1. Science 320, 1496–1501. doi:10.1126/science.1157535.

14. Sancak, Y., Bar-Peled, L., Zoncu, R., Markhard, A.L., Nada, S., and Sabatini, D.M. (2010). Ragulator-Rag complex targets mTORC1 to the lysosomal surface and is necessary for its activation by amino acids. Cell 141, 290–303. 10.1016/j.cell.2010.02.024.

15. Kim, E., Goraksha-Hicks, P., Li, L., Neufeld, T.P., and Guan, K.-L. (2008). Regulation of TORC1 by Rag GTPases in nutrient response. Nature Cell Biology 10, 935–945. 10.1038/ncb1753.

16. Efeyan, A., Zoncu, R., Chang, S., Gumper, I., Snitkin, H., Wolfson, R.L., Kirak, O., Sabatini, D.D., and Sabatini, D.M. (2013). Regulation of mTORC1 by the Rag GTPases is necessary for neonatal autophagy and survival. Nature 493, 679–683. 10.1038/nature11745.

17. Bar-Peled, L., Schweitzer, L.D., Zoncu, R., and Sabatini, D.M. (2012). Ragulator is a GEF for the rag GTPases that signal amino acid levels to mTORC1. Cell 150, 1196–1208. 10.1016/j.cell.2012.07.032.

18. Binda, M., Péli-Gulli, M.-P., Bonfils, G., Panchaud, N., Urban, J., Sturgill, T.W., Loewith, R., and De Virgilio, C. (2009). The Vam6 GEF Controls TORC1 by Activating the EGO Complex. Molecular Cell 35, 563–573. 10.1016/j.molcel.2009.06.033.

19. Dubouloz, F., Deloche, O., Wanke, V., Cameroni, E., and De Virgilio, C. (2005). The TOR and EGO Protein Complexes Orchestrate Microautophagy in Yeast. Molecular Cell 19, 15–26. 10.1016/j.molcel.2005.05.020.

20. Bar-Peled, L., Chantranupong, L., Cherniack, A.D., Chen, W.W., Ofna, K.A., Grabiner, B.C., Spear, E.D., Carter, S.L., Meyerson, M., and Sabatini, D.M. (2013). A Tumor Suppressor Complex with GAP Activity for the Rag GTPases That Signal Amino Acid Sufficiency to mTORC1. Science 340, 1100–1106. doi:10.1126/science.1232044.

21. Yang, H., Jiang, X., Li, B., Yang, H.J., Miller, M., Yang, A., Dhar, A., and Pavletich, N.P. (2017). Mechanisms of mTORC1 activation by RHEB and inhibition by PRAS40. Nature 552, 368–373. 10.1038/nature25023.

22. Menon, S., Dibble, C.C., Talbott, G., Hoxhaj, G., Valvezan, A.J., Takahashi, H., Cantley, L.C., and Manning, B.D. (2014). Spatial control of the TSC complex integrates insulin and nutrient regulation of mTORC1 at the lysosome. Cell 156, 771–785. 10.1016/j.cell.2013.11.049.

23. Dibble, C.C., and Manning, B.D. (2013). Signal integration by mTORC1 coordinates nutrient input with biosynthetic output. Nature Cell Biology 15, 555–564. 10.1038/ncb2763.

24. Inoki, K., Zhu, T., and Guan, K.-L. (2003). TSC2 Mediates Cellular Energy Response to Control Cell Growth and Survival. Cell 115, 577–590. 10.1016/S0092-8674(03)00929-2.

25. Inoki, K., Li, Y., Xu, T., and Guan, K.-L. (2003). Rheb GTPase is a direct target of TSC2 GAP activity and regulates mTOR signaling. Genes & Development 17, 1829–1834. 10.1101/gad.1110003.

26. Gwinn, D.M., Shackelford, D.B., Egan, D.F., Mihaylova, M.M., Mery, A., Vasquez, D.S., Turk, B.E., and Shaw, R.J. (2008). AMPK Phosphorylation of Raptor Mediates a Metabolic Checkpoint. Molecular Cell 30, 214–226. 10.1016/j.molcel.2008.03.003.

27. Yuan, H.-X., Wang, Z., Yu, F.-X., Li, F., Russell, R.C., Jewell, J.L., and Guan, K.-L. (2015). NLK phosphorylates Raptor to mediate stress-induced mTORC1 inhibition. Genes & Development 29, 2362–2376. 10.1101/gad.265116.115.

28. Jewell, J.L., Fu, V., Hong, A.W., Yu, F.-X., Meng, D., Melick, C.H., Wang, H., Lam, W.-L.M., Yuan, H.-X., Taylor, S.S., and Guan, K.-L. (2019). GPCR signaling inhibits mTORC1 via PKA phosphorylation of Raptor. eLife 8, e43038. 10.7554/eLife.43038.

29. Meng, D., Yang, Q., Melick, C.H., Park, B.C., Hsieh, T.-S., Curukovic, A., Jeong, M.-H., Zhang, J., James, N.G., and Jewell, J.L. (2021). ArfGAP1 inhibits mTORC1 lysosomal localization and activation. The EMBO Journal 40, e106412. 10.15252/embj.2020106412.

30. Li, L., Kim, E., Yuan, H., Inoki, K., Goraksha-Hicks, P., Schiesher, R.L., Neufeld, T.P., and Guan, K.-L. (2010). Regulation of mTORC1 by the Rab and Arf GTPases *. Journal of Biological Chemistry 285, 19705–19709. 10.1074/jbc.C110.102483.

31. Panchaud, N., Péli-Gulli, M.-P., and De Virgilio, C. (2013). Amino Acid Deprivation Inhibits TORC1 Through a GTPase-Activating Protein Complex for the Rag Family GTPase Gtr1. Science Signaling 6, ra42–ra42. 10.1126/scisignal.2004112.

32. Neklesa, T.K., and Davis, R.W. (2009). A Genome-Wide Screen for Regulators of TORC1 in Response to Amino Acid Starvation Reveals a Conserved Npr2/3 Complex. PLOS Genetics 5, e1000515. 10.1371/journal.pgen.1000515.

33. Laxman, S., Sutter, B.M., Shi, L., and Tu, B.P. (2014). Npr2 inhibits TORC1 to prevent inappropriate utilization of glutamine for biosynthesis of nitrogen-containing metabolites. Science Signaling 7, ra120–ra120. 10.1126/scisignal.2005948.

34. Chen, J., Sutter, B.M., Shi, L., and Tu, B.P. (2017). GATOR1 regulates nitrogenic cataplerotic reactions of the mitochondrial TCA cycle. Nature Chemical Biology 13, 1179–1186. 10.1038/nchembio.2478.

35. Algret, R., Fernandez-Martinez, J., Shi, Y., Kim, S.J., Pellarin, R., Cimermancic, P., Cochet, E., Sali, A., Chait, B.T., Rout, M.P., and Dokudovskaya, S. (2014). Molecular Architecture and Function of the SEA Complex, a Modulator of the TORC1 Pathway *. Molecular & Cellular Proteomics 13, 2855–2870. 10.1074/mcp.M114.039388.

36. Hatakeyama, R. (2021). Pib2 as an Emerging Master Regulator of Yeast TORC1. Biomolecules 11, 1489.

37. Kim, A., and Cunningham, K.W. (2015). A LAPF/phafin1-like protein regulates TORC1 and lysosomal membrane permeabilization in response to endoplasmic reticulum membrane stress. Molecular Biology of the Cell 26, 4631–4645. 10.1091/mbc.E15-08-0581.

38. Troutman, K.K., Varlakhanova, N.V., Tornabene, B.A., Ramachandran, R., and Ford, M.G.J. (2022). Conserved Pib2 regions have distinct roles in TORC1 regulation at the vacuole. Journal of Cell Science 135. 10.1242/jcs.259994.

39. Tarassov, K., Messier, V., Landry, C.R., Radinovic, S., Molina, M.M.S., Shames, I., Malitskaya, Y., Vogel, J., Bussey, H., and Michnick, S.W. (2008). An in Vivo Map of the Yeast Protein Interactome. Science 320, 1465–1470. doi:10.1126/science.1153878.

40. Tanigawa, M., and Maeda, T. (2017). An In Vitro TORC1 Kinase Assay That Recapitulates the Gtr-Independent Glutamine-Responsive TORC1 Activation Mechanism on Yeast Vacuoles. LID - 10.1128/MCB.00075-17 [doi] LID - e00075–17.

41. Michel, A.H., Hatakeyama, R., Kimmig, P., Arter, M., Peter, M., Matos, J., De Virgilio, C., and Kornmann, B. (2017). Functional mapping of yeast genomes by saturated transposition. eLife 6, e23570. 10.7554/eLife.23570.

42. Ukai, H., Araki, Y., Kira, S., Oikawa, Y., May, A.I., and Noda, T. (2018). Gtr/Ego-independent TORC1 activation is achieved through a glutamine-sensitive interaction with Pib2 on the vacuolar membrane. PLOS Genetics 14, e1007334. 10.1371/journal.pgen.1007334.

43. Tanigawa, M., Yamamoto, K., Nagatoishi, S., Nagata, K., Noshiro, D., Noda, N.N., Tsumoto, K., and Maeda, T. (2021). A glutamine sensor that directly activates TORC1. Communications Biology 4, 1093. 10.1038/s42003-021-02625-w.

44. Varlakhanova, N.V., Mihalevic, M.J., Bernstein, K.A., and Ford, M.A.-O. (2017). Pib2 and the EGO complex are both required for activation of TORC1.

45. Brito, A.S., Soto Diaz, S., Van Vooren, P., Godard, P., Marini, A.M., and Boeckstaens, M. (2019). Pib2-Dependent Feedback Control of the TORC1 Signaling Network by the Npr1 Kinase. iScience 20, 415–433. 10.1016/j.isci.2019.09.025.

46. Yerlikaya, S., Meusburger, M., Kumari, R., Huber, A., Anrather, D., Costanzo, M., Boone, C., Ammerer, G., Baranov, P.V., and Loewith, R. (2016). TORC1 and TORC2 work together to regulate ribosomal protein S6 phosphorylation in Saccharomyces cerevisiae. Mol Biol Cell 27, 397–409. 10.1091/mbc.E15-08-0594.

47. Chen, X., Wang, G., Zhang, Y., Dayhoff-Brannigan, M., Diny, N.L., Zhao, M., He, G., Sing, C.N., Metz, K.A., Stolp, Z.D., et al. (2018). Whi2 is a conserved negative regulator of TORC1 in response to low amino acids. PLoS Genet 14, e1007592. 10.1371/journal.pgen.1007592.

48. Hughes Hallett, J.E., Luo, X., and Capaldi, A.P. (2014). State Transitions in the TORC1 Signaling Pathway and Information Processing in Saccharomyces cerevisiae. Genetics 198, 773–786. 10.1534/genetics.114.168369.

49. Soulard, A., Cremonesi, A., Moes, S., Schütz, F., Jenö, P., and Hall, M.N. (2010). The Rapamycin-sensitive Phosphoproteome Reveals That TOR Controls Protein Kinase A Toward Some But Not All Substrates. Molecular Biology of the Cell 21, 3475–3486. 10.1091/mbc.e10-03-0182.

50. Shin, C.S., Kim, S.Y., and Huh, W.K. (2009). TORC1 controls degradation of the transcription factor Stp1, a key effector of the SPS amino-acid-sensing pathway in Saccharomyces cerevisiae. J Cell Sci 122, 2089–2099. 10.1242/jcs.047191.

51. Dokladal, L., Stumpe, M., Hu, Z., Jaquenoud, M., Dengjel, J., and De Virgilio, C. (2021). Phosphoproteomic responses of TORC1 target kinases reveal discrete and convergent mechanisms that orchestrate the quiescence program in yeast. Cell Rep 37, 110149. 10.1016/j.celrep.2021.110149.

52. Cherkasova, V.A., and Hinnebusch, A.G. (2003). Translational control by TOR and TAP42 through dephosphorylation of eIF2alpha kinase GCN2. Genes Dev 17, 859–872. 10.1101/gad.1069003.

53. Kamada, Y., Yoshino, K., Kondo, C., Kawamata, T., Oshiro, N., Yonezawa, K., and Ohsumi, Y. (2010). Tor directly controls the Atg1 kinase complex to regulate autophagy. Mol Cell Biol 30, 1049–1058. 10.1128/MCB.01344-09.

54. Urban, J., Soulard, A., Huber, A., Lippman, S., Mukhopadhyay, D., Deloche, O., Wanke, V., Anrather, D., Ammerer, G., Riezman, H., et al. (2007). Sch9 is a major target of TORC1 in Saccharomyces cerevisiae. Mol Cell 26, 663–674. 10.1016/j.molcel.2007.04.020.

55. Huber, A., French, S.L., Tekotte, H., Yerlikaya, S., Stahl, M., Perepelkina, M.P., Tyers, M., Rougemont, J., Beyer, A.L., and Loewith, R. (2011). Sch9 regulates ribosome biogenesis via Stb3, Dot6 and Tod6 and the histone deacetylase complex RPD3L. EMBO J 30, 3052–3064. 10.1038/emboj.2011.221.

56. Albers, E., Laize, V., Blomberg, A., Hohmann, S., and Gustafsson, L. (2003). Ser3p (Yer081wp) and Ser33p (Yil074cp) are phosphoglycerate dehydrogenases in Saccharomyces cerevisiae. J Biol Chem 278, 10264–10272. 10.1074/jbc.M211692200.

57. Grauslund, M., Didion, T., Kielland-Brandt, M.C., and Andersen, H.A. (1995). BAP2, a gene encoding a permease for branched-chain amino acids in Saccharomyces cerevisiae. Biochim Biophys Acta 1269, 275–280. 10.1016/0167-4889(95)00138-8.

58. Schreve, J.L., Sin, J.K., and Garrett, J.M. (1998). The Saccharomyces cerevisiae YCC5 (YCL025c) gene encodes an amino acid permease, Agp1, which transports asparagine and glutamine. J Bacteriol 180, 2556–2559. 10.1128/JB.180.9.2556-2559.1998.

59. Tanaka, J., and Fink, G.R. (1985). The histidine permease gene (HIP1) of Saccharomyces cerevisiae. Gene 38, 205–214. 10.1016/0378-1119(85)90219-7.

60. Zhu, X., Garrett, J., Schreve, J., and Michaeli, T. (1996). GNP1, the high-affinity glutamine permease of S. cerevisiae. Curr Genet 30, 107–114. 10.1007/s002940050108.

61. MacDiarmid, C.W., Gaither, L.A., and Eide, D. (2000). Zinc transporters that regulate vacuolar zinc storage in Saccharomyces cerevisiae. EMBO J 19, 2845–2855. 10.1093/emboj/19.12.2845.

62. Zhao, H., and Eide, D.J. (1997). Zap1p, a metalloregulatory protein involved in zinc-responsive transcriptional regulation in Saccharomyces cerevisiae. Mol Cell Biol 17, 5044–5052. 10.1128/MCB.17.9.5044.

63. Lewis, D.A., and Bisson, L.F. (1991). The HXT1 gene product of Saccharomyces cerevisiae is a new member of the family of hexose transporters. Mol Cell Biol 11, 3804–3813. 10.1128/mcb.11.7.3804-3813.1991.

64. Boeckstaens, M., Merhi, A., Llinares, E., Van Vooren, P., Springael, J.Y., Wintjens, R., and Marini, A.M. (2015). Identification of a Novel Regulatory Mechanism of Nutrient Transport Controlled by TORC1-Npr1-Amu1/Par32. PLoS Genet 11, e1005382. 10.1371/journal.pgen.1005382.

65. MacGurn, J.A., Hsu, P.C., Smolka, M.B., and Emr, S.D. (2011). TORC1 regulates endocytosis via Npr1-mediated phosphoinhibition of a ubiquitin ligase adaptor. Cell 147, 1104–1117. 10.1016/j.cell.2011.09.054.

66. Schmidt, A., Beck, T., Koller, A., Kunz, J., and Hall, M.N. (1998). The TOR nutrient signalling pathway phosphorylates NPR1 and inhibits turnover of the tryptophan permease. EMBO J 17, 6924–6931. 10.1093/emboj/17.23.6924.

67. Schmidt, A., Hall, M.N., and Koller, A. (1994). Two FK506 resistance-conferring genes in Saccharomyces cerevisiae, TAT1 and TAT2, encode amino acid permeases mediating tyrosine and tryptophan uptake. Mol Cell Biol 14, 6597–6606. 10.1128/mcb.14.10.6597-6606.1994.

68. Worley, J., Luo, X., and Capaldi, A.P. (2013). Inositol pyrophosphates regulate cell growth and the environmental stress response by activating the HDAC Rpd3L. Cell Rep 3, 1476–1482. 10.1016/j.celrep.2013.03.043.

69. Sorger, P.K., and Pelham, H.R. (1988). Yeast heat shock factor is an essential DNA-binding protein that exhibits temperature-dependent phosphorylation. Cell 54, 855–864. 10.1016/s0092-8674(88)91219-6.

70. Chang, Y., and Huh, W.K. (2018). Ksp1-dependent phosphorylation of eIF4G modulates post-transcriptional regulation of specific mRNAs under glucose deprivation conditions. Nucleic Acids Res 46, 3047–3060. 10.1093/nar/gky097.

71. Paczia, N., Becker-Kettern, J., Conrotte, J.F., Cifuente, J.O., Guerin, M.E., and Linster, C.L. (2019). 3-Phosphoglycerate Transhydrogenation Instead of Dehydrogenation Alleviates the Redox State Dependency of Yeast de Novo l-Serine Synthesis. Biochemistry 58, 259–275. 10.1021/acs.biochem.8b00990.

72. Kinoshita, E., Yamada, A., Takeda, H., Kinoshita-Kikuta, E., and Koike, T. (2005). Novel immobilized zinc(II) affinity chromatography for phosphopeptides and phosphorylated proteins. J Sep Sci 28, 155–162. 10.1002/jssc.200401833.

73. Wallace, R.L., Lu, E., Luo, X., and Capaldi, A.P. (2022). Ait1 regulates TORC1 signaling and localization in budding yeast. eLife 11, e68773. 10.7554/eLife.68773.

74. Stracka, D., Jozefczuk, S., Rudroff, F., Sauer, U., and Hall, M.N. (2014). Nitrogen source activates TOR (target of rapamycin) complex 1 via glutamine and independently of Gtr/Rag proteins. J Biol Chem 289, 25010–25020. 10.1074/jbc.M114.574335.

75. Marion, R.M., Regev, A., Segal, E., Barash, Y., Koller, D., Friedman, N., and O’Shea, E.K. (2004). Sfp1 is a stress- and nutrient-sensitive regulator of ribosomal protein gene expression. Proceedings of the National Academy of Sciences 101, 14315–14322. doi:10.1073/pnas.0405353101.

76. Lempiäinen, H., Uotila, A., Urban, J., Dohnal, I., Ammerer, G., Loewith, R., and Shore, D. (2009). Sfp1 Interaction with TORC1 and Mrs6 Reveals Feedback Regulation on TOR Signaling. Molecular Cell 33, 704–716. 10.1016/j.molcel.2009.01.034.

77. Lippman, S.I., and Broach, J.R. (2009). Protein kinase A and TORC1 activate genes for ribosomal biogenesis by inactivating repressors encoded by *Dot6* and its homolog *Tod6*. Proceedings of the National Academy of Sciences 106, 19928–19933. doi:10.1073/pnas.0907027106.

78. Huber, A., French Sl Fau - Tekotte, H., Tekotte H Fau - Yerlikaya, S., Yerlikaya S Fau - Stahl, M., Stahl M Fau - Perepelkina, M.P., Perepelkina Mp Fau - Tyers, M., Tyers M Fau - Rougemont, J., Rougemont J Fau - Beyer, A.L., Beyer Al Fau - Loewith, R., and Loewith, R. (2011). Sch9 regulates ribosome biogenesis via Stb3, Dot6 and Tod6 and the histone deacetylase complex RPD3L.

79. Liko, D., Slattery, M.G., and Heideman, W. (2007). Stb3 Binds to Ribosomal RNA Processing Element Motifs That Control Transcriptional Responses to Growth in <em>Saccharomyces cerevisiae</em>*. Journal of Biological Chemistry 282, 26623–26628. 10.1074/jbc.M704762200.

80. Bianchi, F., Van’t Klooster, J.S., Ruiz, S.J., and Poolman, B. (2019). Regulation of Amino Acid Transport in Saccharomyces cerevisiae. Microbiol Mol Biol Rev 83. 10.1128/MMBR.00024-19.

81. De Craene, J.O., Soetens, O., and Andre, B. (2001). The Npr1 kinase controls biosynthetic and endocytic sorting of the yeast Gap1 permease. J Biol Chem 276, 43939–43948. 10.1074/jbc.M102944200.

82. Cui, Z., Napolitano, G., de Araujo, M.E.G., Esposito, A., Monfregola, J., Huber, L.A., Ballabio, A., and Hurley, J.H. (2023). Structure of the lysosomal mTORC1-TFEB-Rag-Ragulator megacomplex. Nature 614, 572–579. 10.1038/s41586-022-05652-7.

83. Napolitano, G., Di Malta, C., Esposito, A., de Araujo, M.E.G., Pece, S., Bertalot, G., Matarese, M., Benedef, V., Zampelli, A., Stasyk, T., et al. (2020). A substrate-specific mTORC1 pathway underlies Birt-Hogg-Dube syndrome. Nature 585, 597–602. 10.1038/s41586-020-2444-0.

84. Mattaini, K.R., Sullivan, M.R., and Vander Heiden, M.G. (2016). The importance of serine metabolism in cancer. J Cell Biol 214, 249–257. 10.1083/jcb.201604085.

85. Storici, F., and Resnick, M.A. (2006). The delitto perfetto approach to in vivo site-directed mutagenesis and chromosome rearrangements with synthetic oligonucleotides in yeast.

86. Storici, F., Lewis, L.K., and Resnick, M.A. (2001). In vivo site-directed mutagenesis using oligonucleotides. Nature Biotechnology 19, 773–776. 10.1038/90837.

87. Mülleder, M., Campbell, K., Matsarskaia, O., Eckerstorfer, F., and Ralser, M. (2016). Saccharomyces cerevisiae single-copy plasmids for auxotrophy compensation, multiple marker selection, and for designing metabolically cooperating communities.

88. Urban, J., Soulard A Fau - Huber, A., Huber A Fau - Lippman, S., Lippman S Fau - Mukhopadhyay, D., Mukhopadhyay D Fau - Deloche, O., Deloche O Fau - Wanke, V., Wanke V Fau - Anrather, D., Anrather D Fau - Ammerer, G., Ammerer G Fau - Riezman, H., Riezman H Fau - Broach, J.R., et al. (2007). Sch9 is a major target of TORC1 in Saccharomyces cerevisiae.

89. Licklider, L.J., Thoreen, C.C., Peng, J., and Gygi, S.P. (2002). Automation of Nanoscale Microcapillary Liquid Chromatography−Tandem Mass Spectrometry with a Vented Column. Analytical Chemistry 74, 3076–3083. 10.1021/ac025529o.

90. Taus, T., Köcher, T., Pichler, P., Paschke, C., Schmidt, A., Henrich, C., and Mechtler, K. (2011). Universal and Confident Phosphorylation Site Localization Using phosphoRS. Journal of Proteome Research 10, 5354–5362. 10.1021/pr200611n.

91. Schindelin, J., Arganda-Carreras, I., Frise, E., Kaynig, V., Longair, M., Pietzsch, T., Preibisch, S., Rueden, C., Saalfeld, S., Schmid, B., et al. (2012). Fiji: an open-source plazorm for biological-image analysis. Nature Methods 9, 676–682. 10.1038/nmeth.2019.

