## Supplementary material for "The Molecular Logic of Gtr1/2 and Pib2 Dependent TORC1 Regulation in Budding Yeast": Table S2

| Dataset | Ser 20 | Ser 22 | Ser 27 | Thr 28 | Ser 29 | Thr 31 | Ser 33 |
|---|---|---|---|---|---|---|---|
| Fig 4 (rap dependence) | x | x | | x | x | | |
| Soulard (rap dependence) | x | x | | | x | x | |
| Dokládal (rap dependence) | x | x | x | x | | | x |
| Ser33-IP in SC (rep 1) | x | x | x | | | | x |
| Ser33-IP in rap (rep 1) | | x | | | | | x |
| Difference (rep 1) | x | | x | | | | |
| Ser33-IP in SC (rep 2) | x | x | x | x | x | x | x |
| Ser33-IP in rap (rep 2) | x | x | | | | x | x |
| Difference (rep 2) | x | | x | x | x | | x |

**Table 2. Rapamycin dependent phosphorylation sites in Ser33.** The top three rows show the position of the phosphorylation sites that decrease abundance after rapamycin treatment, as identified in Figure 4 and Table 1 (top row), and in the published work of Soulard (second row[1]) and Dokládal (third row; found in at least 3/5 experiments[2]). The fourth and fifth rows show the phosphosites identified via mass-spectrometry based mapping of Ser33 immunopurified from cells growing in log growth phase in SC medium and one hour after treatment with 200 ng/ml rapamycin (samples were prepared together and examined in back-to-back mass spec runs). The sixth row shows the sites that decrease in abundance comparing the log growth and rapamycin samples. Rows 7-9 show data for a biological replicate of rows 4-6. In second IP a peptide was identified that is phosphorylated at sites 20 and 33 in SC medium (green Xs), but not rapamycin, indicating that the abundance of one, or both, of these sites decreases (orange Xs). Dokládal also found a rapamycin dependent site at Ser 2 that we did not pick up and was not mutated in our study. The raw data for rows 4-9 is in Tables S4-S7.


1. Soulard, A., Cremonesi, A., Moes, S., Schütz, F., Jenö, P., and Hall, M.N. (2010). The Rapamycin-sensitive Phosphoproteome Reveals That TOR Controls Protein Kinase A Toward Some But Not All Substrates. Molecular Biology of the Cell *21*, 3475-3486. 10.1091/mbc.e10-03-0182.
2. Dokladal, L., Stumpe, M., Hu, Z., Jaquenoud, M., Dengjel, J., and De Virgilio, C. (2021). Phosphoproteomic responses of TORC1 target kinases reveal discrete and convergent mechanisms that orchestrate the quiescence program in yeast. Cell Rep *37*, 110149. 10.1016/j.celrep.2021.110149.
